# Phosphorylation of Nephrin induces phase separated domains that move through actomyosin contraction

**DOI:** 10.1101/558965

**Authors:** Soyeon Kim, Joseph M. Kalappurakkal, Satyajit Mayor, Michael K. Rosen

**Author notes:** Correspondence to Michael K. Rosen.

## Abstract

The plasma membrane of eukaryotic cells is organized into lipid and protein microdomains, whose assembly mechanisms and functions are incompletely understood. We demonstrate that proteins in the Nephrin/Nck/N-WASP actin-regulatory pathway cluster into micron-scale domains at the basal plasma membrane upon triggered phosphorylation of transmembrane Nephrin. The domains are persistent but readily exchange components with their surroundings, and their formation is dependent on the number of Nck SH3 domains, suggesting they are phase separated polymers assembled through multivalent interactions among the three proteins. The domains form independent of the actin cytoskeleton, but acto-myosin contractility induces their rapid lateral movement. Nephrin phosphorylation induces larger clusters at the cell periphery, which are associated with extensive actin assembly and dense filopodia. Our studies illustrate how multivalent interactions between proteins at the plasma membrane can produce micron-scale organization of signaling molecules, and how the resulting clusters can both respond to and control the actin cytoskeleton.

## Introduction

The organization of the plasma membrane into biochemically distinct compartments is believed to play an important role in the transmission of signals between the extracellular environment and the cytoplasm (Astro & de Curtis, 2015, Day & Kenworthy, 2009, Edidin, 2003, Grecco, Schmick et al., 2011). Spatially distinct membrane domains ranging in size from a few nanometers to one or two micrometers have been observed in many biological settings and during many cellular processes (Kusumi, Suzuki et al., 2011, van Zanten & Mayor, 2015). Prominently among these, numerous receptors involved in cell growth, neuronal signaling, and cell adhesion form oligomers upon binding to extracellular ligands, and this oligomerization often leads to formation of higher-order clusters (Grecco et al., 2011, Groves & Kuriyan, 2010, Honerkamp-Smith, Veatch et al., 2009, Kusumi et al., 2011). While the initial oligomerization is understood in structural terms in many cases, the physical mechanisms that produce the larger structures in response to signals and the functional consequences of those structures are less well understood.

Many instances of receptor clustering implicate stochastic oligomerization of proteins at the plasma membrane as a key component of microdomain formation (Astro & de Curtis, 2015, Kholodenko, Hoek et al., 2000, Wu, 2013). For example, oligomerization of cadherins requires *trans-*interaction between cadherins presented by the neighboring cells as well as *cis-*interactions (Wu, Vendome et al., 2011). Also the death-inducing signaling complex is formed through homotypic association of death domains (DD) from CD95 and FADD, followed by procaspase activation and initiation of apoptotic pathway (Schleich, Warnken et al., 2012). Interestingly, numerous receptors and their interacting proteins contain multivalent domains. These receptors have been observed as clustered structures at the plasma membrane (Astro & de Curtis, 2015, Groves & Kuriyan, 2010, Wu, Pan et al.). Examples includes Ephrin receptors with three intracellular phosphorylation sites and LAT with four intracellular phosphorylation sites (Nikolov, Xu et al., 2013, Su, Ditlev et al., 2016, Yap, Gomez et al., 2015).

Interactions between multivalent membrane proteins and their multivalent ligands can drive oligomerization and concomitant phase separation of protein components on lipid bilayers (Banjade & Rosen, 2014, Su et al., 2016). This has been demonstrated biochemically in several analogous systems consisting of a membrane receptor phosphorylated on multiple tyrosine residues, an adaptor protein containing a Src Homology 2 (SH2) domain and multiple SH3 domains, and a third protein containing multiple Proline Rich Motifs (PRMs). Interactions between SH2 and phosphotyrosine (SH2-pTyr) and SH3-PRM interactions between the molecules then drive assembly and concomitant phase separation. Among the cell adhesion signaling molecules in kidney podocytes, phase separation has been observed for Nephrin, Nck and N-WASP (Banjade & Rosen, 2014, Banjade, Wu et al., 2015); and among T cell receptor (TCR) signaling molecules it has been observed for LAT, Grb2 and SOS, and also for LAT, Gads and Slp-76 (Su et al., 2016). In the first two systems the threshold concentration for phase separation *in vitro* was shown to be dependent on the valency of one or more species, and in the Nephrin system it was also dependent on SH2-pTyr affinity. The Nephrin study, as well as a more complex reconstitution of the TCR pathway, also showed that the protein clusters promote local actin polymerization in the presence of the Arp2/3 complex (Banjade & Rosen, 2014, Su et al., 2016). Experiments in Jurkat T cells involving perturbations of the number of pTyr sites in LAT suggested that multivalency-driven phase separation underlies formation of LAT clusters *in vivo* (Su et al., 2016). However, parallel cellular work has not yet been reported on the Nephrin pathway, nor has the importance of valency of other modular interactions been explored *in vivo*.

A notable difference between artificial synthetic bilayers and the crowded cellular environment is the presence of the cortical actomyosin network at the plasma membrane. This network, which is composed of actin filaments cross-linked with myosin II motors, plays critical roles in diverse cell activities that require mechanical transduction, including migration and morphological change (Paluch, Sykes et al., 2006). A number of studies suggest that the actomyosin system also functions to spatially organize signaling cascades through its interactions with various actin binding receptors and adaptors. For example, CD36, a receptor in macrophages, forms clusters through movement within confinement regions defined by the cortical cytoskeleton (Jaqaman & Grinstein, 2012). GPI anchored proteins have also been shown to form nanoclusters through coupling to a dynamically contracting network of short actin filaments and myosin at the plasma membrane (Gowrishankar, Ghosh et al., 2012, Koster, Husain et al., 2016, Koster & Mayor, 2016, Rao & Mayor, 2014). Finally, during T cell activation actin retrograde flow induces radial movement of clusters containing the TCR, LAT and various adaptor proteins at the immunological synapse between a T cell and an antigen promoting cell (APC) (Jaqaman & Grinstein, 2012, Sherman, Barr et al., 2011, Yu, Smoligovets et al., 2013).

Here, we extend our work on multivalency-mediated phase separation of receptors, by showing that micron-scale clusters of Nephrin/Nck/N-WASP can be rapidly induced at the basal plasma membrane of HeLa cells through triggered phosphorylation of Nephrin. Formation of the clusters is dependent on both the cellular concentration and SH3-valency of Nck1. Nephrin, Nck and N-WASP all exchange readily between the clusters and the surrounding medium, as assessed by fluorescence recovery after photobleaching (FRAP), although their dynamics is four times slower within the clusters compared to the regions without clusters. Interestingly, actomyosin contraction promotes rapid movement of the clusters across the plasma membrane, while formation of the clusters is insensitive to perturbations of the actin cytoskeleton. At longer times, Nephrin phosphorylation also generates larger, Nephrin/Nck/N-WASP clusters at the cell periphery, which coincide with dense, actin-rich arrays of filopodia. Unlike the basal clusters, these peripheral clusters are dependent on actin assembly. Together, these observations suggest that even in a cellular environment crowded with competing signaling molecules, multivalent interactions between Nephrin, Nck and N-WASP can promote formation of phase separated polymers analogous to those previously observed on model membranes. In cells, these polymeric clusters can both respond to the actomyosin cytoskeleton, and also generate new actin-based structures and changes in cell morphology. These investigations suggest general mechanisms by which multivalent signaling molecules can be organized on micron length scales and interact with the cortical actin cytoskeleton.

## Results

### Phosphorylation of Nephrin can be induced using FKBP-FRB heterodimers

Nephrin is an essential structural component of the kidney glomerulus, a specialized structure for ultrafiltration of plasma, consisting of eight extracellular IgG repeats, a transmembrane domain, and an unstructured tail of ∼160 amino acids (Martin & Jones, 2018, Ruotsalainen, Ljungberg et al., 1999). Nephrin maintains extracellular junctions between foot processes of podocyte cells in the kidney, while also regulating actin cytoskeletal support within the cells (Martin & Jones, 2018, Welsh & Saleem, 2010). At these interfaces, the eight extracellular IgG domains of Nephrin interact in-trans between cells to create an inter-digitating pattern of tight junctions that act as a biological filter (Gerke, 2003). Electron tomographic imaging of fixed kidney tissues revealed a network of winding Nephrin strands with a length of 35 nm (Wartiovaara, Ofverstedt et al., 2004). A recent study employing cryo-electron tomography suggested that Nephrin forms a layered assembly structure at the apical region of foot processes between murine podocytes (Grahammer, Wigge et al., 2016).

Phosphorylation of tyrosine residues on the intracellular tail of Nephrin at the filtration barrier is mainly dependent on the Src-family kinase, Fyn (Verma, Wharram et al., 2003). It remains unclear what upstream signals activate Fyn *in vivo*. Fyn can be transiently activated through antibody engagement of the Nephrin extracellular domains, interactions thought to mimic trans-cellular engagement of the Nephrin extracellular domains (Verma, Kovari et al., 2006). However, a recent study suggested that Fyn may be activated by integrin signaling rather than extracellular engagement of Nephrin (Verma, Venkatareddy et al., 2016). Three tyrosine residues are essential for Nephrin signaling to the actin cytoskeleton (Jones, Blasutig et al., 2006). Upon phosphorylation, the Src homology 2 (SH2)/SH3 adaptor protein Nck binds these sites through its SH2 domain (Jones, New et al., 2009). Interestingly, Nck also works in a positive-feedback mechanism, promoting activation of Fyn upon binding to Nephrin (New, Keyvani Chahi et al., 2013). The three SH3 domains of Nck can interact with proline-rich motifs of N-WASP, an activator of the actin nucleation factor, the Arp2/3 complex (Blasutig, New et al., 2008, Buday, Wunderlich et al., 2002, Padrick & Rosen, 2010). Thus, through this pathway, sites of Nephrin phosphorylation recruit and activate the Arp2/3 complex to induce local actin assembly (Buday et al., 2002).

To test whether our *in vitro* observations of p-Nephrin clustering in the presence of Nck and N-WASP (Banjade & Rosen, 2014, Banjade et al., 2015) could be recapitulated *in vivo*, we sought a means of controlling phosphorylation of Nephrin in cells. Previous studies showed that when Nephrin is crosslinked with an extracellular ligand and secondary antibody, its cytoplasmic tail becomes phosphorylated and intracellular signaling is activated (Jones, 2006; Blaustig, 2008). However, this extracellular crosslinking itself is sufficient to cause substantial Nephrin clustering in the absence of a signaling-competent intracellular domain (Jones, 2006; Blaustig, 2008). We sought a purely intracellular means of triggering Nephrin phosphorylation. We employed a system based on the proteins FKBP and FRB, which bind each other with high affinity only in the presence of the small molecule, rapamycin (Banaszynski, Liu et al., 2005). As illustrated in Figure 1A, we used this system to target a Src family kinase to membrane-bound Nephrin upon addition of rapamycin, thus triggering phosphorylation of Nephrin in response to the small molecule cue. To isolate the functions of the Nephrin intracellular tail, we utilized a construct containing the N-terminal signal sequence, transmembrane domain and cytoplasmic domain of the protein, but lacking the large extracellular Ig-repeat element. This Nephrin fragment was fused C-terminally to FRB followed by a fluorescent protein (mEGFP or mCherry), with (GlyGlySer)_n_ linkers between each domain (called Nephrin-FRB hereafter).

**Figure 1.**
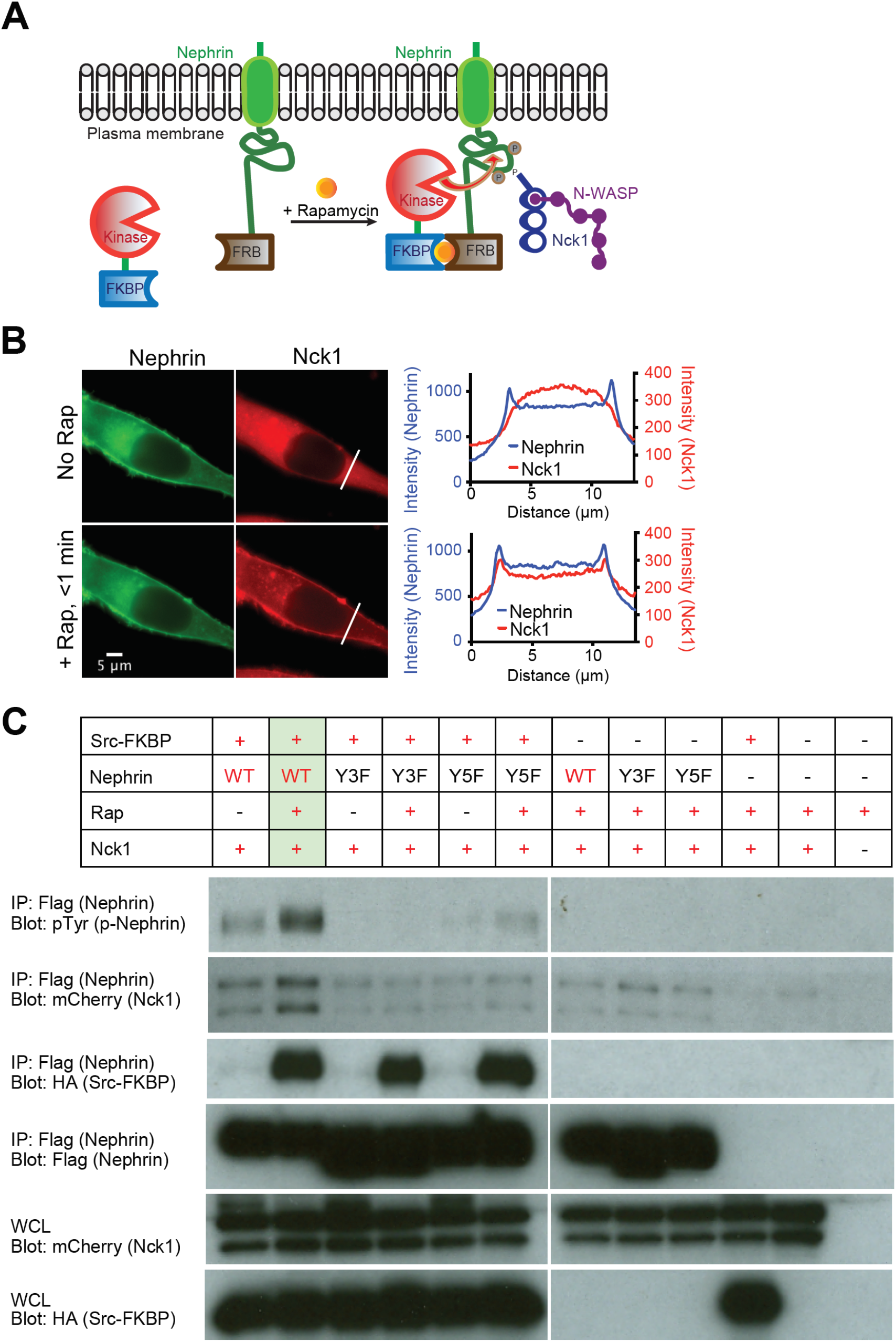
A new tool to manipulate the phosphorylation state of Nephrin in cells. (A) Cartoon illustrating the interactions between Src-FKBP and Nephrin-FRB at the plasma membrane upon rapamycin treatment. Without rapamycin Nephrin-FRB localizes to the plasma membrane while Src-FKBP resides in cytoplasm (left). Rapamycin addition recruits Src-FKBP to Nephrin-FRB, resulting phosphorylation of Nephrin at three sites followed by interactions with Nck1/N-WASP (right). (B) Epi fluorescence images of a live HeLa cell expressing Src-FKBP (not shown), Nephrin-FRB (left column, green) and Nck1 (right column, red) before (top row) or 1 min after (bottom row) addition of rapamycin to the media. Graphs show intensity profiles of indicated lines from the images of Nephrin (blue) and Nck1 (red) on the same row. (C) Western blots of HeLa cells expressing subsets of Src-FKBP, Nephrin(WT or Y3F as indicated)-FRB and Nck1. Antibodies were against pTyr (p-Nephrin), mCherry (Nck1), HA (Src-FKBP), Flag (Nephrin), as indicated. Either whole cell lysates (bottom two rows) or samples immunoprecipitated with anti-Flag affinity gel (top four rows) were used. Cell samples were treated with either DMSO (lanes 1, 3, 5) or rapamycin for 15 min before being lysed.

We co-expressed Nephrin-FRB with a construct containing the catalytic domain of the cSrc tyrosine kinase fused at its C-terminus to FKBP (Src-FKBP hereafter). cSrc is a close relative of the Fyn kinase, which naturally phosphorylates Nephrin in podocytes, and was used as a substitute for Fyn in previous studies (Verma et al., 2003). Addition of rapamycin to cells expressing both Src-FKBP and membrane-associated Nephrin-FRB should cause close approximation of the two proteins at the membrane, promoting high level phosphorylation of Nephrin tyrosines. The kinase domain of Src has a relatively high K_M_ for its substrates (∼100µM; (Foda, Shan et al., 2015)), and the full length protein is often targeted to substrates through its N-terminal SH2 and SH3 domains (Martin, 2001). Thus, by using the isolated kinase domain we hoped to minimize background phosphorylation of Nephrin prior to rapamycin addition, and to reduce undesired phosphorylation of other cellular targets throughout the experiments.

We transiently co-expressed Nephrin-FRB, Src-FKBP and mEGFP-tagged Nck1 (Nck1 hereafter) in HeLa cells. In the absence of rapamycin, Nephrin-FRB localized to the plasma membrane, while Nck1 remained cytoplasmic (Figure 1B; Supplemental Figure S1A). Upon rapamycin addition to the media, Nck1 rapidly (< 1 minute) translocated to the plasma membrane. Translocation coincided temporally with increased phosphorylation of immunoprecipitated Nephrin observed by anti-pTyr western blotting (Figure 1B; Supplemental Figure S1B). Increased phosphorylation was specific for Nephrin-FRB, according to anti-pTyr western blotting of whole cell extracts (Supplemental Figure S1C). Nck1 translocation was not observed when a non-phosphorylatable mutant form of Nephrin-FRB was used (Nephrin(Y3F), lacking the three known sites of Nephrin tyrosine phosphorylation (Jones et al., 2006), or when kinase was not co-expressed (Supplemental Figure S1D). Together, these data indicate that Nck1 translocation was due to binding to tyrosine-phosphorylated Nephrin-FRB (pNephrin-FRB). Pull-down experiments confirmed the interaction between Nck1 and Nephrin-FRB, as more Nck1 bound Nephrin-FRB following rapamycin treatment (Figure 1C). Paralleling translocation, Nck1 binding was abolished when non-phosphorylatable mutant forms of Nephrin-FRB, Nephrin(Y3F) and Nephrin (Y5F) (two additional possible tyrosine phosphorylation sites were mutated to phenylalanine), were used, or when kinase was not co-expressed (Figure 1C). Similar western blotting data were obtained when the system was engineered in 293 cells (Supplemental Figures S1, C and E). This newly developed experimental platform allows us to observe the real-time behavior of Nephrin and its phosphorylation-dependent interactions with binding partners in cells.

### Nephrin/Nck clusters at the basal plasma membrane in response to rapamycin

To evaluate the consequences of Nephrin-Nck1 interactions at the basal plasma membrane, we analyzed live HeLa cells transiently transfected with Src-FKBP, Nephrin-FRB and Nck1 using total internal reflection fluorescence (TIRF) microscopy. Within 10 minutes of rapamycin addition, Nephrin assembled into clusters on the plasma membrane (Figure 2A and Movie S1). These clusters did not stain with markers for focal adhesions (paxillin) or the endocytic machinery (caveolin or clathrin) indicating that they are independent of these structures (Supplemental Figures S2, A and B). Further, all clustering experiments were performed with concurrent methyl-beta-cyclodextrin (MβCD) treatment to block endocytosis of Nephrin-FRB upon phosphorylation (Qin, Tsukaguchi et al., 2009). Treating cells with MβCD not only helped to increase the Nephrin concentration on the membrane (not shown), but also eliminated the possibility that the clusters are generated primarily by cholesterol-dependent lipid raft formation. Approximately 60% of the Nephrin-FRB clusters were also enriched in Nck1, and we limit our analyses hereafter to these Nephrin-Nck1 structures below (Figure 2A; Supplemental Figure S2C). Similar to the clusters of Nephrin/Nck/N-WASP observed *in vitro* on supported lipid bilayers (Banjade & Rosen, 2014), larger Nephrin-FRB/Nck1 clusters in cells were irregularly-shaped, as observed by both diffraction-limited TIRF microscopy and super-resolution stochastic optical reconstruction microscopy (STORM) (Figure 2B). Significantly fewer clusters were observed in cells expressing Nephrin(Y3F) or Nephrin lacking its two additional cytoplasmic tyrosine residues (Nephrin(Y5F)), or in cells expressing wild type Nephrin-FRB in the absence of rapamycin addition or Src-FKBP co-expression, supporting the idea that cluster formation is largely dependent on interactions between pNephrin-FRB and Nck1 (Figure 2C).

**Figure 2.**
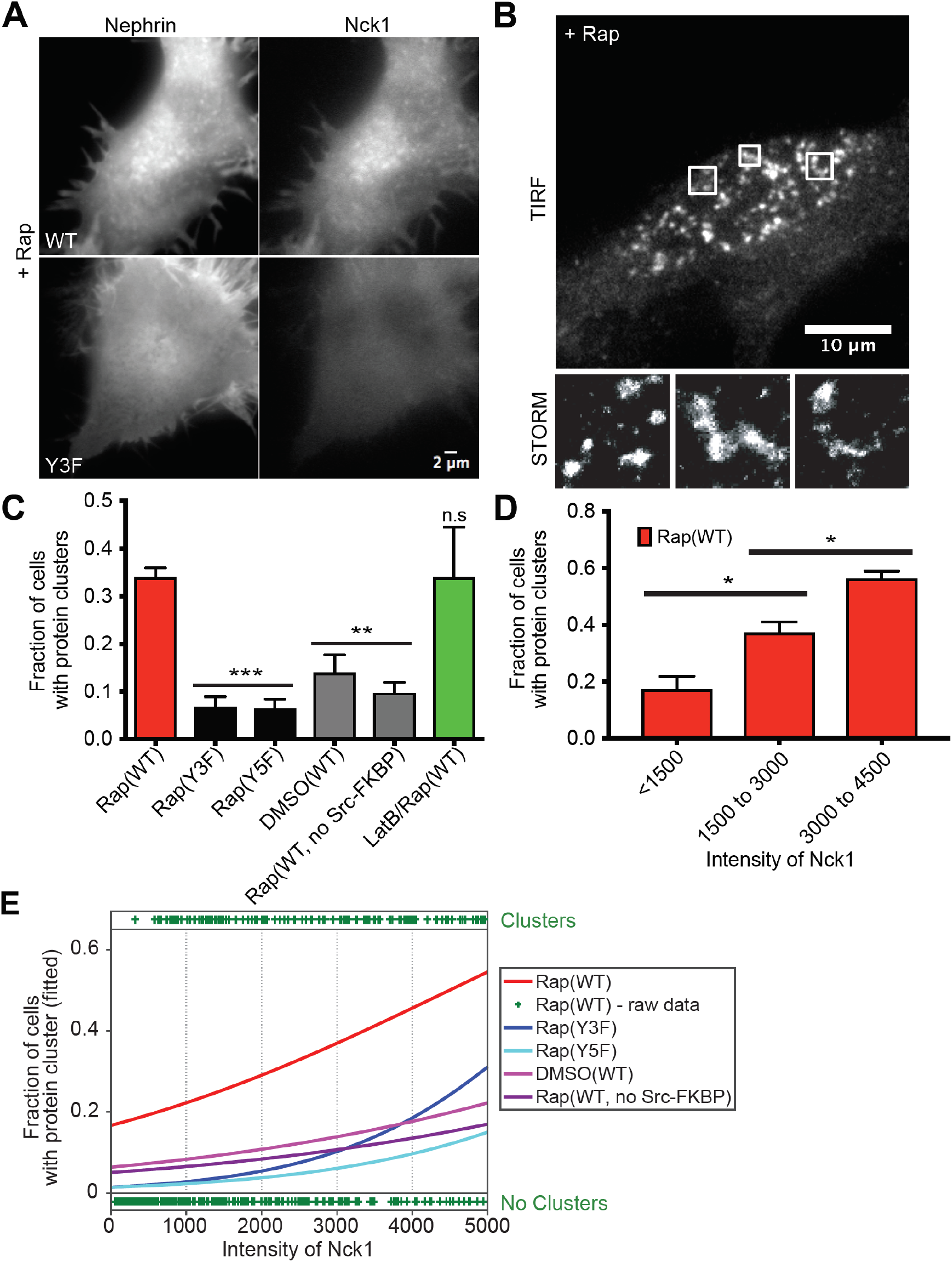
Nephrin/Nck1 cluster formation at the basal membrane is dependent on interactions between Nephrin and Nck1 as well as cellular concentration of Nck1. (A) TIRF images of live HeLa cells expressing Src-FKBP, Nephrin-FRB (left column), and Nck1 (right column). Cells expressed either Nephrin(WT)-FRB (top row) or Nephrin(Y3F)-FRB (bottom row). Both images were taken from cells treated with 10 min of MβCD followed by 30 min of rapamycin. (B) TIRF-STORM images of a fixed HeLa cell expressing Src-FKBP, Nephrin-FRB (shown in images), and Nck1. Cells were fixed after 10 min of MβCD and 30 min of rapamycin treatment. Zoomed images from three boxed regions of the top image are shown in the bottom row. (C to E) Quantitative analyses of protein clustering in various experimental conditions and at different cellular concentrations of Nck1. TIRF images of Nephrin and epi/TIRF images of Nck1 from fixed HeLa cells were used for the analyses. Cells expressed Src-FKBP (unless noted as ‘no Src-FKBP’), Nck1 and Nephrin(WT, Y3F, or Y5F)-FRB as indicated. Cells were treated with 10 min of MβCD and either 30 min of rapamycin (Rap), 30 min of DMSO (DMSO), or 10 min of latrunculin B followed by 20 min of rapamycin (LatB/Rap) (N = 3 independent experiments, each with > 140 cell images). For C and D, error bars represent s.e.m. (*p < 0.05; **p < 0.01; ***p < 0.001) (C) Graph showing the fraction of cells with clusters over various experimental conditions. (D) Graph showing the fraction of cells with clusters for different cytoplasmic expression levels (assessed by epi fluorescence intensity measurements) of Nck1. (E) Graphs showing the fitted probability of a cell forming clusters at a given Nck1 fluorescence intensity level in the cytoplasm. Curves show data for each condition - binary cell responses (y = 1 or 0 for a cell containing or lacking clusters, respectively) and Nck1 intensity values - fit with a logistic regression to estimate the probability of cells producing clusters at a given Nck1 intensity level (lines, left axis). Raw data from the ‘Rap(WT)’ sample are shown as an example with plus signs at the top or bottom of the image, indicating a single cell containing or lacking clusters, respectively, at the indicated Nck1 intensity.

Because multivalency-driven phase separation is dependent on the concentrations of the interacting species (Banjade & Rosen, 2014, Li, Banjade et al., 2012), we examined whether protein clustering depends on cellular concentrations of Nephrin-FRB or Nck1. We used the natural variation in transient transfection to produce a wide range of protein expression levels in HeLa cells. We binned the cells according to the average fluorescence intensity of Nephrin on the basal membrane or Nck1 in the cytoplasm, and determined the probability of observing clusters following rapamycin treatment in each bin. Statistical comparison of the bins shows a significant increase in cluster probability with increasing Nck concentration (Figure 2D; Supplemental Figure S2D). We note that here and below, the appearance of clusters with increasing Nck1 is much less sharp than in our previous *in vitro* experiments (Banjade & Rosen, 2014). This difference is likely due to the fact that we have not controlled for variable N-WASP concentrations in our cellular experiments, and moreover that pNephrin and Nck1 have numerous additional binding partners in cells that can contribute to or compete with phase separation. When averaged across a population, these variables would tend to broaden the concentration dependence of phase separation, even when the transition is sharp in any given cell.

To better compare the different expression constructs and cell treatments, we fit the data using logistic regression (Figure 2E), a model often used to study the association between a binary dependent variable and continuous independent variables. The dependent variable here has a value of 1 or 0 if a cell contains or lacks protein clusters, respectively (+ symbol top or bottom, respectively, in Figure 2E. The independent variables are Nephrin or Nck concentrations. Fitting the data with a logistic regression model returned the probability function of a cell producing protein clusters for variable Nephrin or Nck concentrations. This analysis showed a clear distinction between experimental and control samples (cells expressing Nephrin(Y3F), cells expressing Nephrin(Y5F), cells without Src-FKBP expression, cells without rapamycin treatment), with measured p-values < 0.0001 (Figure 2E; Supplemental Figure S2E). Similar analyses showed that within the ranges observed here, cluster formation was insensitive to Nephrin density (Supplemental Figures S2, E-G).

### Nephrin/Nck1 clusters form independently of actin polymerization

As described above, the cortical actin cytoskeleton is known to contribute to the organization of membrane receptors in several systems. Here we observed that Nephrin/Nck1 clusters produced after rapamycin treatment remained on the plasma membrane when cells were treated later with Latrunculin B (LatB) (Supplemental Figure S3A). To determine whether the cytoskeleton can affect the initial formation of Nephrin/Nck1 clusters, we inhibited actin assembly by treatment with LatB for 10 minutes prior to inducing Nephrin-FRB phosphorylation with rapamycin. This pre-treatment with LatB changed the shape of cells and cortical actin structures but did not obviously alter the organization of Nephrin-FRB, which remained relatively uniformly distributed across the plasma membrane (Figure 3A). Subsequent addition of rapamycin increased phosphorylation of Nephrin-FRB (Supplemental Figure S3B) and induced recruitment of Nck1 to the plasma membrane within one minute, followed by the formation of micron-sized clusters over the ensuing 30 minutes (Figure 3B and Movie S2). Cluster formation was significantly enhanced when cells contained pNephrin-FRB and Nck1 as in experiments above without LatB treatment (Supplemental Figure S3C). The percentage of cells showing clusters increased with increasing expression of Nck1 and remained at a similar level with increasing Nephrin-FRB expression (Figures 3, C and D; Supplemental Figures S3, C-F). These observations indicate that clusters of Nephrin/Nck1 at the plasma membrane form independently of actin polymerization.

**Figure 3.**
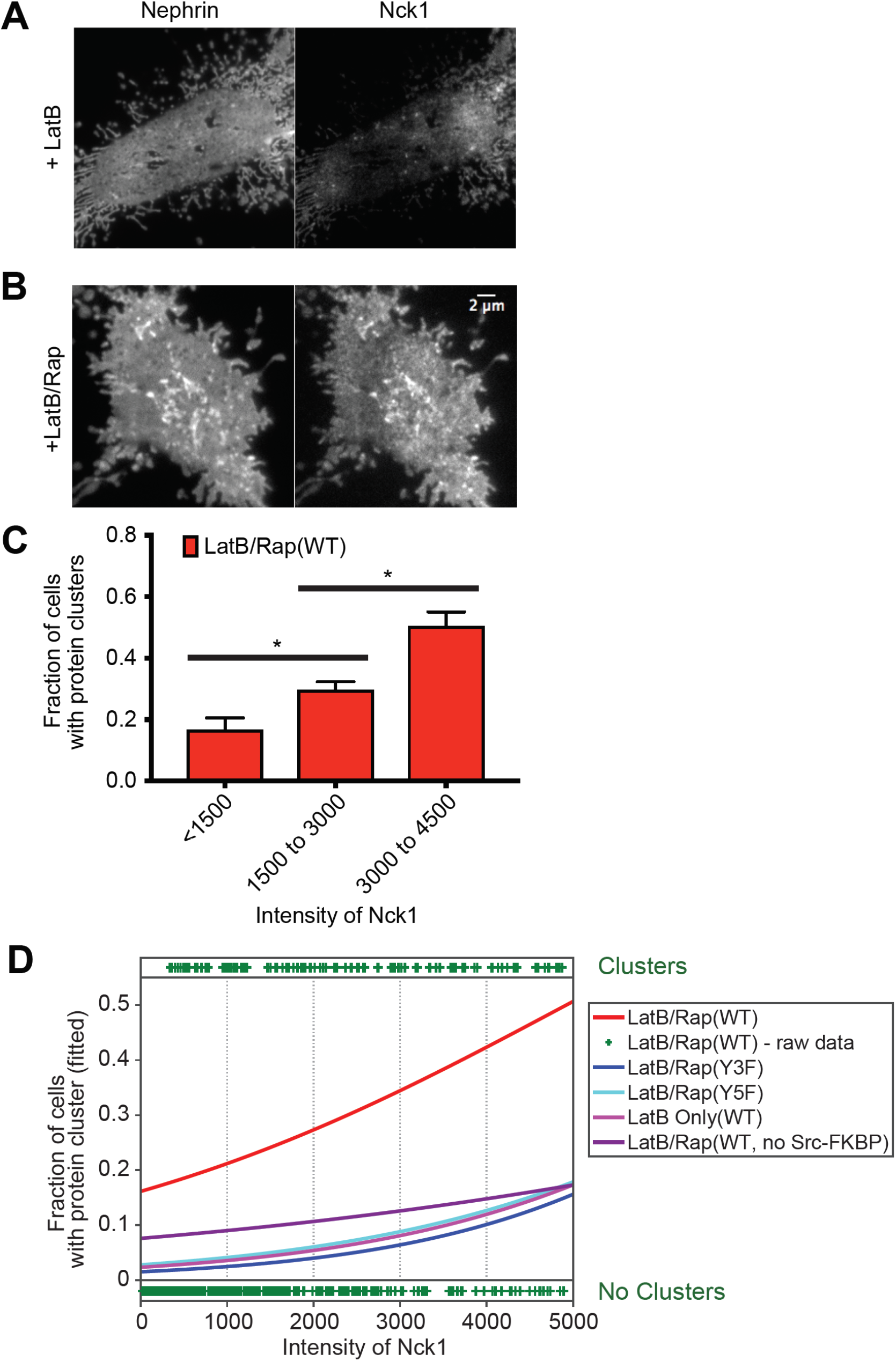
Nephrin/Nck1 clusters form independently of actin polymerization. (A and B) TIRF images of a fixed HeLa cell expressing Src-FKBP (not shown), Nephrin-FRB (left), and Nck1 (right). (A) Cells were treated for 10 min of MβCD followed by 30 min with latrunculin B prior to fixation. (B) Cells were treated for 10 min of MβCD and 15 min of latrunculin B followed by 15 min of rapamycin prior to fixation. (C and D) Quantitative analyses of protein clustering versus cellular concentration of Nck1. TIRF images of Nephrin and epi/TIRF images of Nck1 from fixed HeLa cells were used for the analysis. Cells expressed Src-FKBP (unless noted as ‘no Src-FKBP’), Nck1, and Nephrin(WT, Y3F, or Y5F as indicated)-FRB. Data labeled ‘LatB Only’ were from cells treated with 10 min of MβCD followed by 30 min of latrunculin. All other data (LatB/Rap) are from cells treated with 10 min of MβCD, 15 min of latrunculin B and 15 min of rapamycin prior to fixation. (N = 3 independent experiments, each with > 140 cell images) (C) Graph showing the fraction of cells with clusters for different cytoplasmic expression levels (assessed by epi fluorescence intensity measurements) of Nck1. Error bars, s.e.m. (*p < 0.05) (D) Graphs showing the fitted probability of a cell forming clusters at a given Nck1 intensity level in cytoplasm. Cell data for each condition were fitted (lines) with a logistic regression model (left axis). Each plus sign indicates the status (containing or lacking clusters) of a single cell from ‘LatB/Rap(WT)’ sample (right axis).

### Nephrin clusters have properties consistent with multivalent phase separated polymers

*In vitro* studies have shown that clustering of Nephrin on model membranes and phase separation of the protein in solution are both driven through multivalency-based hetero-oligomerization with Nck1 and N-WASP (Li et al., 2012). A hallmark of this behavior is the dependence of phase separation on the valency of the interacting species. As described above, Nck1 contains three consecutive SH3 domains, which interact with up to six proline-rich motifs in N-WASP (Banjade et al., 2015). To test how Nck1 valency affects protein clustering, we generated a series of proteins composed of tandem repeats of the second SH3 domain of the protein (to eliminate differences in binding affinity of the three natural SH3 domain (Banjade et al., 2015) followed by the SH2 domain. The SH3 domains were connected to each other by the linker that naturally joins the first and second SH3 domains of Nck1 (Figure 4A). These engineered constructs are denoted below as S3, S2, and S1 for proteins with three, two, and one SH3 domain(s), respectively.

**Figure 4.**
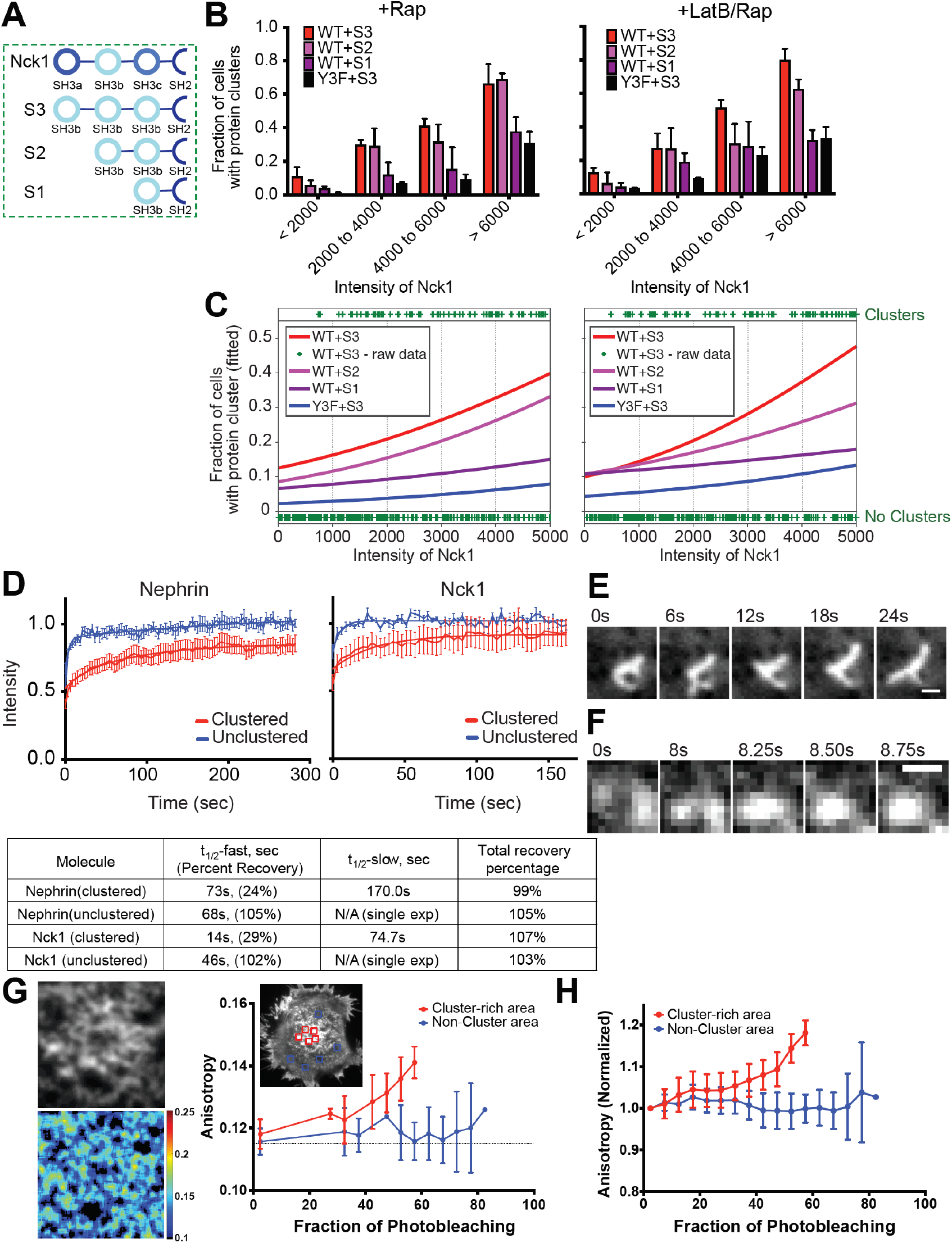
Nephrin/Nck1 clusters at the basal membrane are phase separated structures. (A) Cartoon illustrating wild-type (top) and engineered Nck1 constructs (bottom three) with different SH3 valency. Engineered Nck1 with one, two or three repeats of the second SH3 domain of native Nck1 were named S1, S2, and S3, respectively. The linker connecting the first and second SH3 domains of native Nck1 was used for all inter-SH3 linkers in the engineered proteins. The sequences after the final SH3 domain are identical in all constructs. (B and C) Quantitative analyses of protein clustering for different valencies and cellular concentration of Nck proteins. TIRF images of Nephrin and epi/TIRF images of Nck proteins from fixed HeLa cells were used for the analysis. Cells expressed Src-FKBP (unless noted as ‘no Src-FKBP’), Nephrin(WT or Y3F as indicated)-FRB, and one of the engineered Nck constructs, S3, S2 or S1. Graphs on the left were from cells treated with 10 min of MβCD followed by 30 min of rapamycin (+ Rap), while those on the right were from cells treated with 10 min of MβCD and 15 min of latrunculin B followed by 15 min of rapamycin (+ LatB/Rap) prior to fixation. (N = 3 independent experiments, each with > 125 cell images) (B) Graph showing the fraction of cells with clusters for different cytoplasmic expression levels (determined by epi fluorescence intensity) of Nck1. Error bars, s.e.m. (C) Graphs showing the probability of a cell forming clusters at a given engineered Nck1 construct intensity level in cytoplasm. Cell data for each condition were fitted (lines) with a logistic regression model (left axis). Each plus sign indicates the status (containing or lacking clusters) of a single cell from WT + S3 in each condition (right axis). (D) Fluorescence recovery after photobleaching (FRAP) of Nephrin (left) and Nck1 (right) at the basal membrane in either clustered regions (red) or unclustered regions (blue). FRAP experiments were performed with live cells expressing Src-FKBP, Nephrin-FRB, and Nck1. Cell samples were treated with 10 min of MβCD and 15 min of latrunculin B followed by 15 min of rapamycin prior to imaging. F-tests were performed to distinguish between single or double exponential decay functions. Bars, s.e.m. N ≥ 7 cells. Fitted parameters are shown in the bottom table. (E and F) Timecourse images of clusters live HeLa cells expressing Src-FKBP, Nephrin-FRB (shown in images) and Nck1. Images were acquired every 250 msec for 30 sec. Prior to imaging, cells were treated with 10 min of MβCD and 30 min of rapamycin (Rap). (E) Distinctive morphological changes of a cluster occur within a time interval of 6 sec. (F) Merging of two clusters within a time interval of < 1 sec. (G) Homo-FRET between Nephrin-FRB molecules in a cell shown by fluorescence anisotropy. TIRF images from live HeLa cells expressing Src-FKBP, Nephrin-FRB (shown in the images), and Nck1 were used for the analysis. Left two images are expanded fluorescence (upper) and anisotropy map (bottom) images for a cluster-rich area, selected from the cell shown in the right image of Nephrin-FRB. Graph shows the increase of anisotropy with the decay of fluorescence intensities as fluorophores are bleached during imaging for the single cell shown. Regions used for measurement are indicated by red colored boxes (cluster-rich area) or blue-colored boxes (non-cluster area). Bars, s.d. N = 5 regions. (H) Graph showing normalized anisotropy versus fraction of photobleaching, averaged across five cells. Bars, s.d. N= 5 cells, each with 2-5 regions.

We quantitatively compared cluster formation in cells expressing either S3, S2, or S1 at different expression levels to determine the effect of valency on cluster formation. Logistic regression analysis of these data clearly showed that cells expressing higher valency constructs were more likely to contain clusters with lower expression of the engineered proteins (Figures 4, B and C; Supplemental Figures S4, A and B). Without LatB treatment, cells expressing S2 were nearly as likely as cells expressing S3 to contain clusters (p = 0.8, Figures 4, B and C (left panels)). But in LatB treated cells, the S2 construct had a significantly lower ability to form clusters than S3 (p < 0.001, Figures 4, B and C (right panels)). Under both conditions, cells expressing S1 showed a lower level of clustering than S2 (p < 0.0001) but higher than negative control samples expressing Nephrin(Y3F) only without LatB treatment (p value < 0.01, Figures 4, B and C). Notably, while higher Nephrin density decreased cluster forming ability in cells expressing S3 and S2 (for rapamycin + LatB treatment), cluster formation in cells with S1 was insensitive with Nephrin density (Supplemental Figures S4, A and B). This behavior is consistent with titrating out of the two-phase regime of the phase diagram when an individual protein is in large excess (Banjade & Rosen, 2014, Flory, 1953), an effect that is expected to occur at lower densities/concentrations when the protein has higher valency. Because of this effect there can be an optimal concentration regime to promote multivalency mediated receptor clustering.

Further evidence of multivalency-based oligomerization *in vitro*, derived from the dynamic properties of Nephrin/Nck/N-WASP clusters formed on supported lipid bilayers (Banjade & Rosen, 2014). Within the clusters, all three molecules showed rapid, bi-phasic recovery in FRAP (fluorescence recovery after photobleaching) experiments. Both recovery phases were appreciably faster for Nck and N-WASP than for Nephrin (t_1/2,fast_ ∼ 1.5 vs 60 sec; t_1/2,slow_ ∼ 40 vs 360 sec). In regions of the bilayer outside the clusters, recovery of Nephrin was mono-exponential and more rapid (t_1/2_ = 21 sec). Further, the macroscopic clusters merged with each other on seconds to minutes timescales. We analogously characterized the dynamics of Nephrin/Nck clusters in cells using FRAP. As in the *in vitro* experiments, the cellular Nephrin/Nck puncta show bi-phasic recovery kinetics on tens to hundreds of seconds (Figure 4D). Recovery is again faster for Nck than for Nephrin (t_1/2,fast_ = 14 and 73 sec, respectively; t_1/2,slow_ = 75 and 170 sec, respectively), likely because Nck recovers largely through exchange with the cytoplasm while Nephrin recovers exclusively through movement within the membrane (Figure 4D). Outside of the clusters, both Nephrin and Nck1 recover with single-exponential kinetics, and more rapidly and to a greater extent than within the clusters (t_1/2_ = 69 and 47 seconds, respectively, Figure 4D; Supplemental Figure S4C). Finally, amorphous large clusters changed shape dynamically on timescales of several seconds, and we occasionally saw smaller clusters encounter each other and merge during imaging. These events had a timecourses of one-to-several seconds. Together, these results support our proposal that the clusters are non-covalent polymer networks that can freely exchange components through the dissociation of SH2-pTyr (and probably SH3-PRM) interactions giving rise to rapid molecular and macroscopic dynamics.

We further evaluated the organization of proteins in the clustered domains by measuring the dependence of the fluorescence emission anisotropy of Nephrin-FRB-YFP on the density of the fluorophore, which reports on homo-fluorescence resonance energy transfer (homo-FRET) between the YFPs (Goswami, Gowrishankar et al., 2008). We found that the YFP fluorescence anisotropy gradually increased as Nephrin clusters were photobleached, indicating homo-FRET between the Nephrin-FRB-YFP molecules (Figures 4, G and H). This result suggests that a significant fraction of Nephrin-FRB-YFP molecules are within FRET-distances (∼ 5 nm for the YFP-YFP pair) nm of each other inside the clusters (Ghosh, Saha et al., 2012), consistent with a network crosslinked by Nck1 and N-WASP. In contrast, outside of clusters YFP fluorescence anisotropy did not change with photobleaching (Figures 4, G and H), indicating that the Nephrin molecules are farther apart in these regions of the cell, suggesting a lack of significant crosslinking.

In the presence of actin and the Arp2/3 complex, Nephrin/Nck/N-WASP clusters potently stimulate the formation of new actin filaments *in vitro* (Banjade & Rosen, 2014). We tried several approaches in both live and fixed cells, but did not observe similar enrichment at the analogous clusters in cells (not shown). However, in all cases background staining of cortical actin was very high, and this may have prevented observation of localized enrichment. Thus, our experimental system does not enable a definitive assessment of this potential activity of the clusters.

Together, these data suggest that our *in vivo* Nephrin clusters are very similar to the phase separated protein structures observed in our previous *in vitro* experiments (Banjade & Rosen, 2014).

### Nephrin clusters move across the plasma membrane through actions of the cortical actomyosin network

Nephrin clusters formed *in vitro* on supported bilayers move very little on the membrane surface (Banjade & Rosen, 2014). Clusters merge over time as their dynamic edges encounter one another, but their centers of mass do not change appreciably. In contrast, the Nephrin clusters formed in cells are highly mobile throughout the time course of our experiments (Figures 5, A (top row) and B; Movie S3). They travel across the plasma membrane, mostly through Brownian like motion although sometimes in linear tracks, with rates as high as 0.5 µm/min. Mean-squared displacement (MSD) analysis revealed that the mean diffusion coefficient of clusters is approximately 6 nm^2^/sec, which is comparable with other clustered receptors, such as nicotinic acetylcholine receptor (nAChR) (Barrantes, Bermudez et al., 2010). This mobility could be decreased by treating cells with LatB either before or after induction of Nephrin phosphorylation by rapamycin (Figures 5, A (second row) and B; Supplemental Figures S5, A and D; Movie S4). Clusters formed in this manner showed only Brownian-like motion on the membrane surface. Blebbistatin, a potent myosin inhibitor, similarly blocked movement of the clusters (Figures 5, A (third row) and B; Movie S5), although its inactive enantiomer had no effect (Supplemental Figures S5, B and D). Small molecule inhibitors of the Arp2/3 complex, CK666 or CK869 (Pollard & Cooper, 2009), also decreased the diffusion coefficient of clusters (Figures 5, A (bottom row) and B; Supplemental Figures S5, C and D; Movie S6). Clusters formed by Nephrin(Y3F), which do not form through Nck1 binding (Supplemental Figure S2C), showed diffusion coefficients similar to that of wild type Nephrin in the presence of Arp2/3 complex inhibitors (Supplemental Figure S5D). It is notable that the MSD of more than 25% of clusters without any drug treatment exhibited power-law exponents (α) larger than 1.0, the expected value from a purely viscous fluid (Supplemental Figure S5E). Thus, cluster mobility on the plasma membrane appears to be dependent on the contractile activity of the cortical actomyosin cytoskeleton as well as the assembly of new filaments by the Arp2/3 complex.

**Figure 5.**
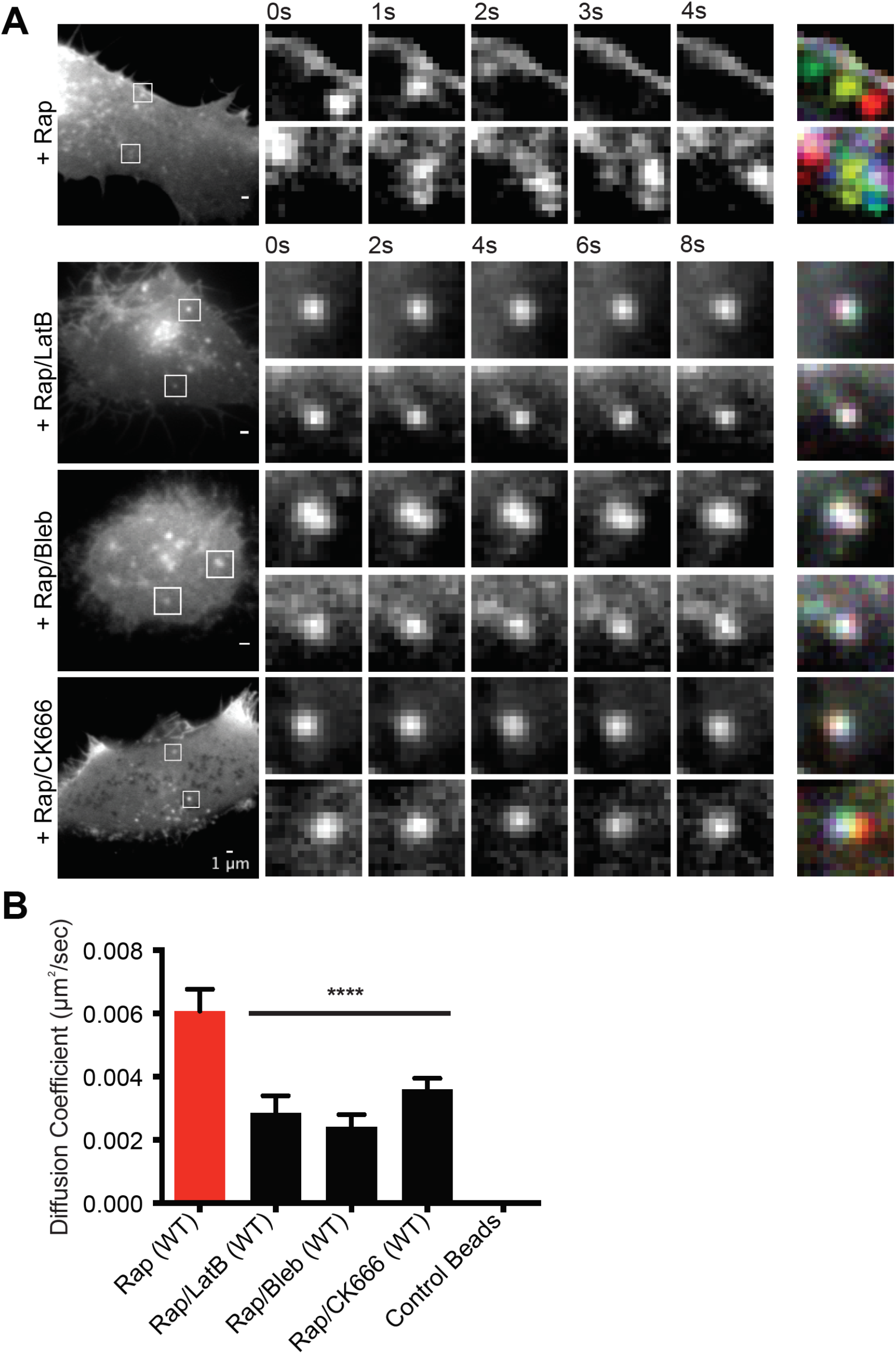
Nephrin clusters move across the plasma membrane through actions of cortical actomyosin. (A, B) TIRF images of live HeLa cells expressing Src-FKBP, Nephrin-FRB (shown in images) and Nck1. Images were recorded every 250 msec for 30 sec. Prior to imaging, cells were treated with 10 min of MβCD and 30 min of rapamycin (Rap) or 10 min of MβCD, 20 min of rapamycin followed by 20 min of either latrunculin B (Rap/LatB), blebbistatin (Rap/Bleb), or CK666 (Rap/CK666). (A) In the left images, two clusters are indicated with arrows in each cell. These clusters are shown in expansion in the time courses to the right. The right-most column shows overlays of the five timecourse images coded with different colors. In this representation, when movement is limited, clusters at different time points superimpose and appear white. Contrast and brightness settings for the different cells were individually optimized to aid visualization of clusters. (B) Calculated diffusion coefficient from the time-lapse images of cells treated with various actomyosin inhibitors. Error bars, s.e.m. N ≥ 14 cells for each condition. (****p < 0.0001)

### Clustering Nephrin with Nck1 promotes peripheral protrusions with a dense actin meshwork

In addition to examining clustering of Nephrin on the basal plasma membrane using TIRF microscopy, we also examined the behavior of Nephrin at the cell periphery using confocal microscopy. Live cell confocal imaging showed a dramatic change in the distribution of Nephrin as well as the morphology of cell edges over a 30 minute time course following rapamycin treatment (Figure 6A). Within ten to twenty minutes of Nck recruitment to the peripheral membrane, cells formed peripheral micron-sized clusters enriched with Nephrin and Nck1. Concurrent with enrichment of the signaling proteins, these clusters generated multiple filopodia-like or microspike structures extending outward from the cell periphery. Staining of fixed cells with Alexa647-phalloidin showed that these morphological changes occurred concomitant with a large increase in F-actin at the sites of Nephrin clustering. These peripheral clusters, observed in the middle confocal section of a cell, where membrane did not attach to the glass surface, showed dependencies analogous to those observed by TIRF imaging of the basal cell surface. Their formation required binding and localization of Nck1 to phosphorylated Nephrin, (Figure 6B). Also the fraction of cells with clusters increased with increasing number of Nck1 SH3 domains (Figure 6C). Notably, when cells were treated initially with rapamycin to induce peripheral clusters, and then LatB secondarily, the clusters persisted for more than 30 minutes along with strong staining for actin filaments (Supplemental Figure S6A).

**Figure 6.**
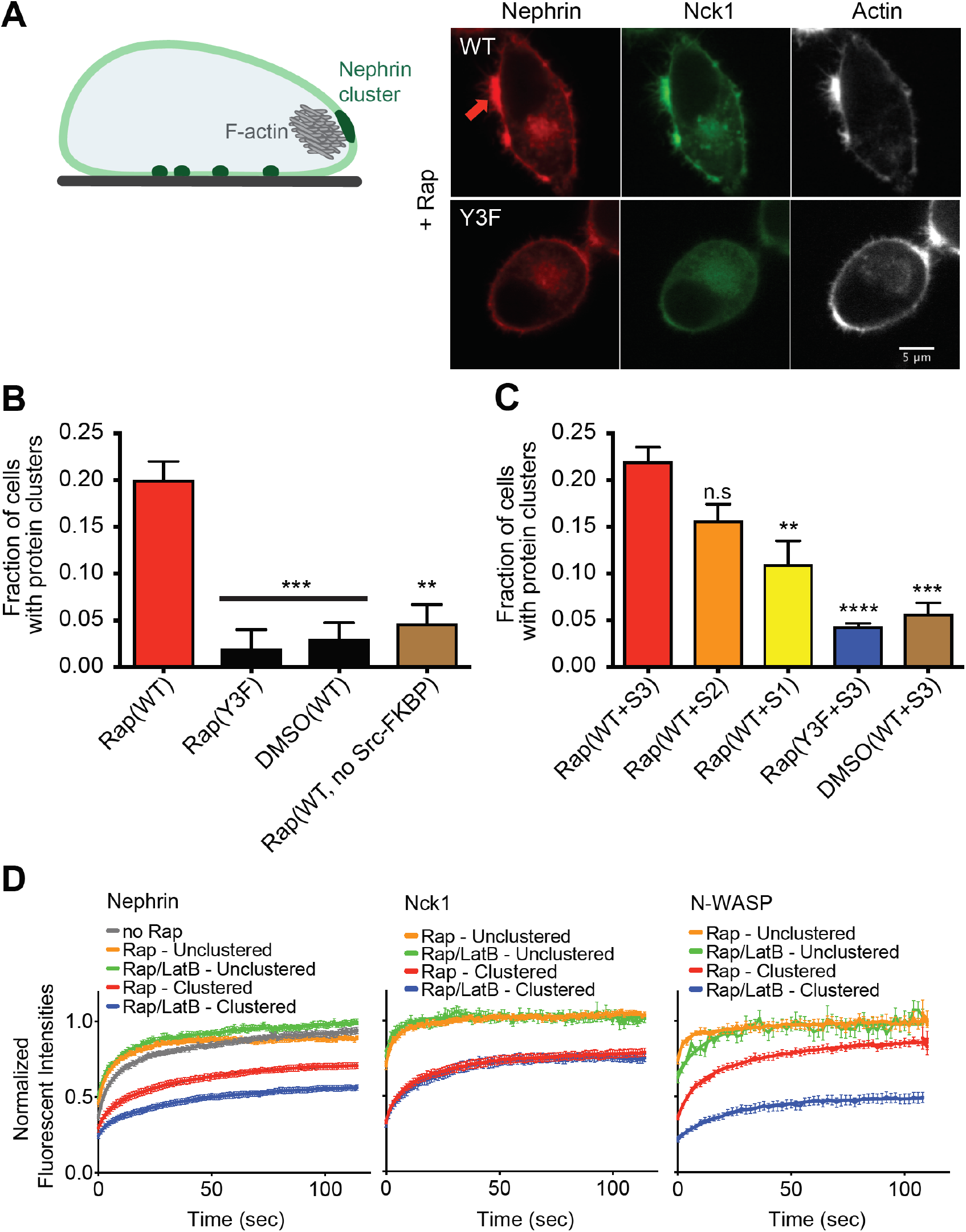
Nephrin/Nck1 clusters at the periphery of cells form with dense actin structures. (A) Left: cartoon illustrating a HeLa cell with Nephrin clusters at the basal membrane (dark green regions at bottom of the cell) and at the periphery (large dark green region at the cell side) enriched with F-actin structure (gray mesh). Right: spinning disk confocal images from fixed Hela cells expressing Src-FKBP, Nephrin(WT, top row; Y3F, bottom row)-FRB (red), and Nck1 (green). Cell samples were treated with 10 min of MβCD followed by 10 min of rapamycin. Fixed cell samples were stained for F-actin using A647-phalloidin (right column, white). (B) Graph showing the fraction of cells with clusters expressing a subset of Src-FKBP (unless noted as ‘no Src-FKBP’), Nephrin(WT or Y3F as indicated)-FRB, and Nck1. Spinning disk confocal images from fixed cell samples were used for the analysis. Cells were treated with 10 min of MβCD followed by either 10 min of rapamycin (Rap) or 10 min of DMSO (DMSO). Error bars, s.e.m. N = 3 independent experiments, each with > 25 cell images. (**p < 0.01, ***p < 0.001) (C) Graph showing the fraction of cells with clusters expressing a subset of Src-FKBP, Nephrin(WT or Y3F as indicated)-FRB, and one of the engineered Nck proteins, S3, S2, or S1. Spinning disk confocal images from fixed cell samples were used for the analysis. Cells were treated with 10 min of MβCD followed by either 10 min of rapamycin (Rap) or 10 min of DMSO (DMSO). Error bars, s.e.m. N = 3 independent experiments, each with > 20 cell images. (**p < 0.01, ***p < 0.001, ****p < 0.0001) (D) Fluorescence recovery after photobleaching (FRAP) of Nephrin (left), Nck1 (middle) or N-WASP (right) at the periphery of a cell in either clustered regions (red and blue) or unclustered region (orange and green). FRAP experiments were performed with live cells expressing Src-FKBP, Nephrin-FRB, and Nck1 with (right) or without (left) N-WASP. Cells were treated with 10 min of MβCD followed by either 10 min of rapamycin (Rap) or 10 min of rapamycin and 10 min of latrunculin B (Rap/LatB). Recovery of Nephrin-FRB, Nck1, or N-WASP intensity in the bleached region was imaged every 1 s for 120 s. Error bars, s.e.m. N > 25 cells analyzed for Nephrin. N ≥ 14 cells analyzed for Nck1 and N-WASP.

Using confocal microscopy, we examined the dynamics of large (2∼3 microns) peripheral Nephrin clusters by FRAP. Similar to the behavior of the basal clusters, Nephrin at the periphery recovered more slowly than Nck1 recovery following photobleaching, and both proteins recovered more slowly within clusters than outside of clusters (Figure 6D; Supplemental Figure S6B). Interestingly, the rate of Nephrin recovery within clusters significantly decreased upon LatB treatment, while Nephrin recovery throughout the rest of membrane was not affected by the compound. This behavior is reminiscent of previous observations that the dynamics of N-WASP at sites of vaccinia virus adhesion to eukaryotic cells are decreased by LatB treatment (Weisswange, Newsome et al., 2009). However, in contrast to vaccinia tails, recovery of Nck1 was unaffected by LatB treatment.

## Discussion

We previously demonstrated that signaling systems based on multiply-phosphorylated Nephrin or LAT, and adaptor proteins consisting of multiple SH2, SH3 and PRM elements, can assembly into micrometer scale clusters *in vitro* on supported lipid bilayers. The sharpness of the transition from homogeneous to clustered receptors, the valency and affinity dependence of the concentration threshold for the transition, and the ready fusion and rapid dynamics of the resulting clusters suggest formation by coupled oligomerization and LLPS. The process is analogous to three dimensional LLPS that has been observed for many multivalent molecular systems (Banani, Rice et al., 2016, Banjade et al., 2015, Fromm, Kamenz et al., 2014, Li et al., 2012, Lin, Protter et al., 2015, Mitrea, Cika et al., 2016, Zeng, Shang et al., 2016), except that the receptor systems are restricted to membranes through attachment of one component to the bilayer. In TCR signaling, we showed a similar dependence of clustering on LAT phosphotyrosine valency and similar fusion behavior of clusters in cells as *in vitro*, suggesting the same process of oligomerization and LLPS drive cluster formation in both environments. Here, we have also seen analogous similarities in cellular and biochemical behaviors of the Nephrin system: valency dependence of the Nck SH3 domains, concentration dependence of cluster formation and dynamic nature of the clusters. Thus, even in the crowded cellular environment, with many potential competitive protein-protein interactions, and natural lipid compositions and lipid interactions, multivalent pTyr-SH2-SH3-PRM interactions are sufficient to promote clustering. This mechanism is likely to be relevant to a variety of other signaling pathways such as Eph/ephrin, cadherin or TCR signaling, which are composed of analogous interactions (Nikolov et al., 2013, Su et al., 2016, Yap et al., 2015).

In TCR signaling, LAT clusters form at the periphery of the interface between the T cell and its conjugated antigen presenting cell (APC), and move radially toward the center of the interface through retrograde flow of the actomyosin cytoskeleton. Similarly vaccinia viruses, which assemble Nck/-N-WASP containing clusters at their membrane-attached surfaces, move across the plasma membrane of infected cells. Here, we have also observed rapid dynamics of Nephrin clusters at the plasma membrane. These are also driven by the actomyosin system. The dependence on the Arp2/3 complex suggests that movement requires assembly of new actin filaments; dependence on myosin motors suggests that movement requires actomyosin contraction. Unlike in T cells, movement of Nephrin clusters is not coordinated, and in this respect is more similar to vaccinia movement. This difference from TCR signaling is probably because our kinase-recruitment approach induces clusters randomly, and because actomyosin in HeLa cells lacks the cooperative movement seen at the T cell-APC interface (Hong, Murugesan et al., 2017, Murugesan, Hong et al., 2016, Yi, Wu et al., 2012). In all three systems the clusters contain Nck and N-WASP (or its homolog, WASP) at some point in their lifetimes. We recently showed that for LAT clusters in T cells, strong binding to actomyosin is mediated through interactions of basic regions in both proteins with actin filaments (Ditlev, Vega et al., 2018). It is likely that similar interactions of these same regions mediate coupling to actomyosin in the Nephrin and vaccinia virus systems as well. Further experiments are necessary to understand whether Nck and N-WASP play similar mechanistic roles in the movement of clusters by dynamic actomyosin in these different but related signaling systems.

The technology we developed here to phosphorylate a receptor on cue could have general utility in studies of signaling. The approach essentially bypasses upstream chemical processes in a signaling pathway (externally-driven receptor clustering, receptor crosstalk, etc.) to activate the output of a single receptor type on a timescale of seconds to minutes. This is achieved simply by fusing FRB to the protein of interest, and co-expressing a Src-FKBP. But utilizing a kinase with high K_M_ (∼low affinity for substrates), we hoped to provide relatively selective phosphorylation of Nephrin over other molecules at the membrane or in the cell. Anti-pTyr western blotting of lysates after rapamycin treatment suggests that this is the case, since p-Nephrin appears to be the dominant phosphorylated species. The approach has weaknesses, including potential phosphorylation of p-Nephrin-associated proteins that are also near the recruited kinase, and the fact that fusion of the receptor to FRB could alter function. Nevertheless, the simplicity of the approach makes it a potentially valuable addition to the toolkit for studying signaling pathways.

It is unclear whether the two types of Nephrin clusters observed here, those on the basal cell surface and those at the cell periphery, are mechanistically related or independent. Both show similar dependencies on the SH3 valency of Nck and in both Nck dynamics are slowed relative to the surrounding membranes. An important difference is that while the basal clusters form independent of actin, the peripheral clusters do not form if the actin network is disrupted prior to rapamycin treatment (although the peripheral Nephrin does appear to be phosphorylated under these conditions, Supplemental Figure S3B, and it recruits Nck). One interesting possibility is that two cluster types are mechanistically related, such that small basal clusters act as precursors, and coalesce into larger clusters at the periphery through actin-based movement. In such a model, disruption of actin with LatB might block peripheral cluster formation through blocking movement, and consequently coalescence, of the basal clusters. Such a model would be consistent with the delayed timing of the peripheral clusters (10-20 minutes post-rapamycin) relative to the basal clusters (5-10 minutes post-rapamycin). Alternatively, the peripheral clusters may form through an independent actin-based process (likely in conjunction with multivalency-based assembly, given their valency dependence and dynamic properties). This could be akin to feedback models proposed for actin-dependent formation of Cdc42 puncta in yeast (Altschuler, Angenent et al., 2008). In these models, initially recruited Cdc42 recruits an actin transport system, which then recruits additional Cdc42. Further studies are needed to understand the mechanism of formation of the peripheral clusters, and their relationship with the basal clusters.

Nephrin is known for its important role in generating the slit diaphragm of the glomerular filtration barrier of the kidney (Kestila, Lenkkeri et al., 1998, Lenkkeri, Mannikko et al., 1999). This structure is formed by extensive cell-cell adhesions along the foot processes of kidney podocytes, epithelial cells that surround the glomerular capillaries (Greka & Mundel, 2012). The extracellular domain of Nephrin is the major structural component of the slit diaphragm (Khoshnoodi, Sigmundsson et al., 2003, Ruotsalainen et al., 1999, Wartiovaara et al., 2004). It has been proposed that during development the initial protrusions from podocytes prior to stable cell-to-cell contact may be mediated by intracellular signaling of Nephrin to the actin cytoskeleton (Greka & Mundel, 2012). For example, the phosphorylation level of Nephrin is increased at these initial stages of foot processes formation, and interactions between Nephrin and Nck are critical for proper nephron formation (Jones et al., 2006, New, Martin et al., 2016, Verma et al., 2006). Nephrin phosphorylation also increases during slit diaphragm recovery after injury, which also requires dynamic changes in cell adhesion (Jones et al., 2009, New et al., 2016, Verma et al., 2006). Mutations of Nephrin or deletion of Nck in the kidney cause major defects in morphology of the slit diaphragm and kidney function (Jones et al., 2006, Jones et al., 2009, Li, Zhu et al., 2006), and a recently developed Nephrin(Y3F)-expressing mouse showed progressive kidney disease and loss of foot processes (New et al., 2016). Notably, out of approximately 150 missense mutation of Nephrin annotated in the HGMD, nine mutations were reported in the cytoplasmic domain and an additional 25 mutations would produce truncations lacking all phosphorylation sites. Further, in multiple disease models, reduced Nephrin phosphorylation levels have been reported (Martin & Jones, 2018). However, it is still largely unknown how Nephrin signaling can result in the formation of the complicated structures seen in the slit diaphragm. Our study proposes that the appearance of actin-rich structures at microdomains condensed with Nephrin may be an initial step for the creation of the interdigitated pattern of podocyte junctions. The perturbations we have generated biochemically and in cells, altering valency and affinity of elements within the Nephrin signaling pathway to produce quantitative changes in clustering, provide potential routes to testing the functional importance of phase separation in kidney development.

## Methods and Materials

### Constructs

All Nephrin-FRB, Src-FKBP were expressed from pcDNA vectors (Invitrogen). The Nephrin-FRB construct was cloned by conjugation of the cytoplasmic tail of Nephrin (residues 1036-1241) and FRB (residues 2015-2114 of human mTOR) tethered by a [GGS]_3_GSGGS linker. The Src-FKBP construct was cloned by conjugation of the kinase domain of human Src (residues 263-522) and FKBP (residues 3-108 of human FKBP12) tethered by a [GGS]_3_ linker. For imaging experiments, mCherry or mEGFP (EGFP with mutation: A206K) was cloned after FKBP or FRB through a (GGS)_3_ linker. All Nephrin-FRB constructs contained a Flag-tag at their C-terminus. All Src-FKBP constructs contained an HA-tag at their C-terminus. Nck1 was cloned into pEGFP vector containing EGFP with A206K mutation (Clontech). Engineered Nck proteins were generated as repeats of the second SH3 domain (SH3b, residues 109-168) of Nck1 connected by the linker that naturally connects the first and second SH3 domains of the protein (residues 62-108) (Banjade et al., 2015). Further information on cloned constructs is provided in Table 1.

**Table 1.**
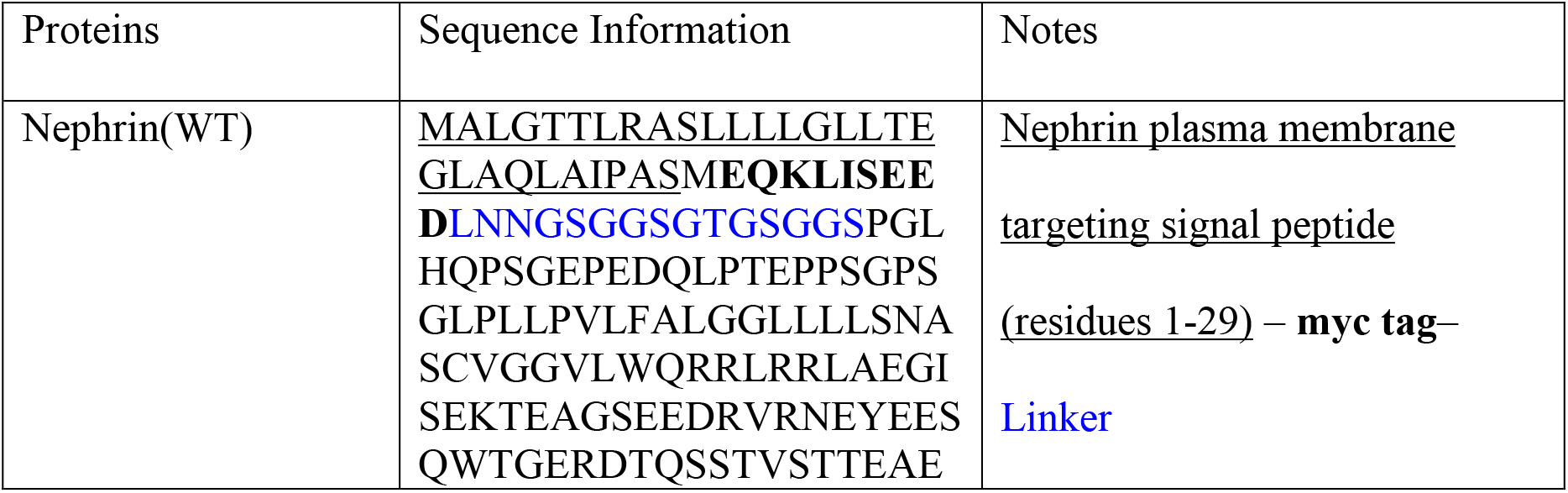
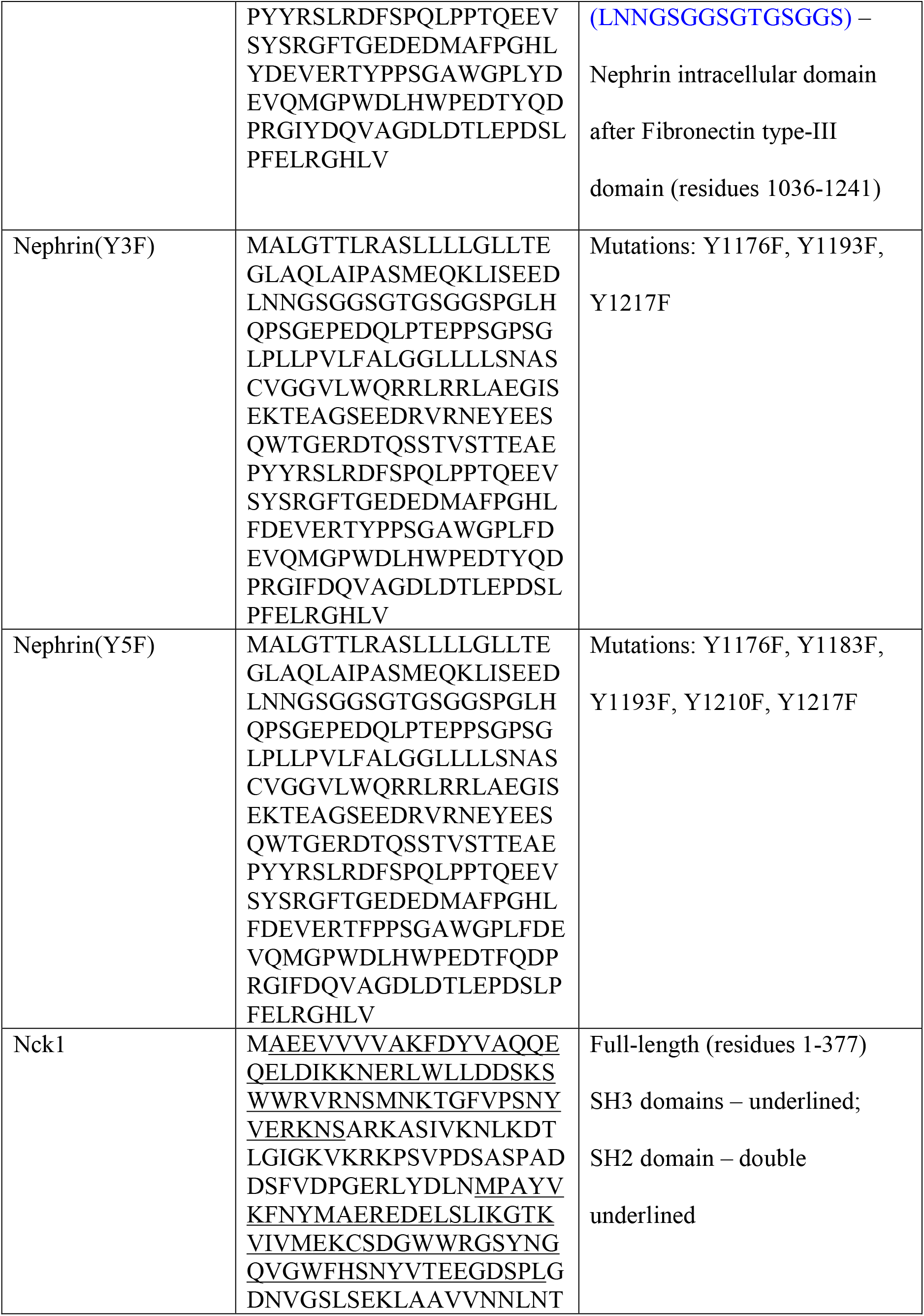
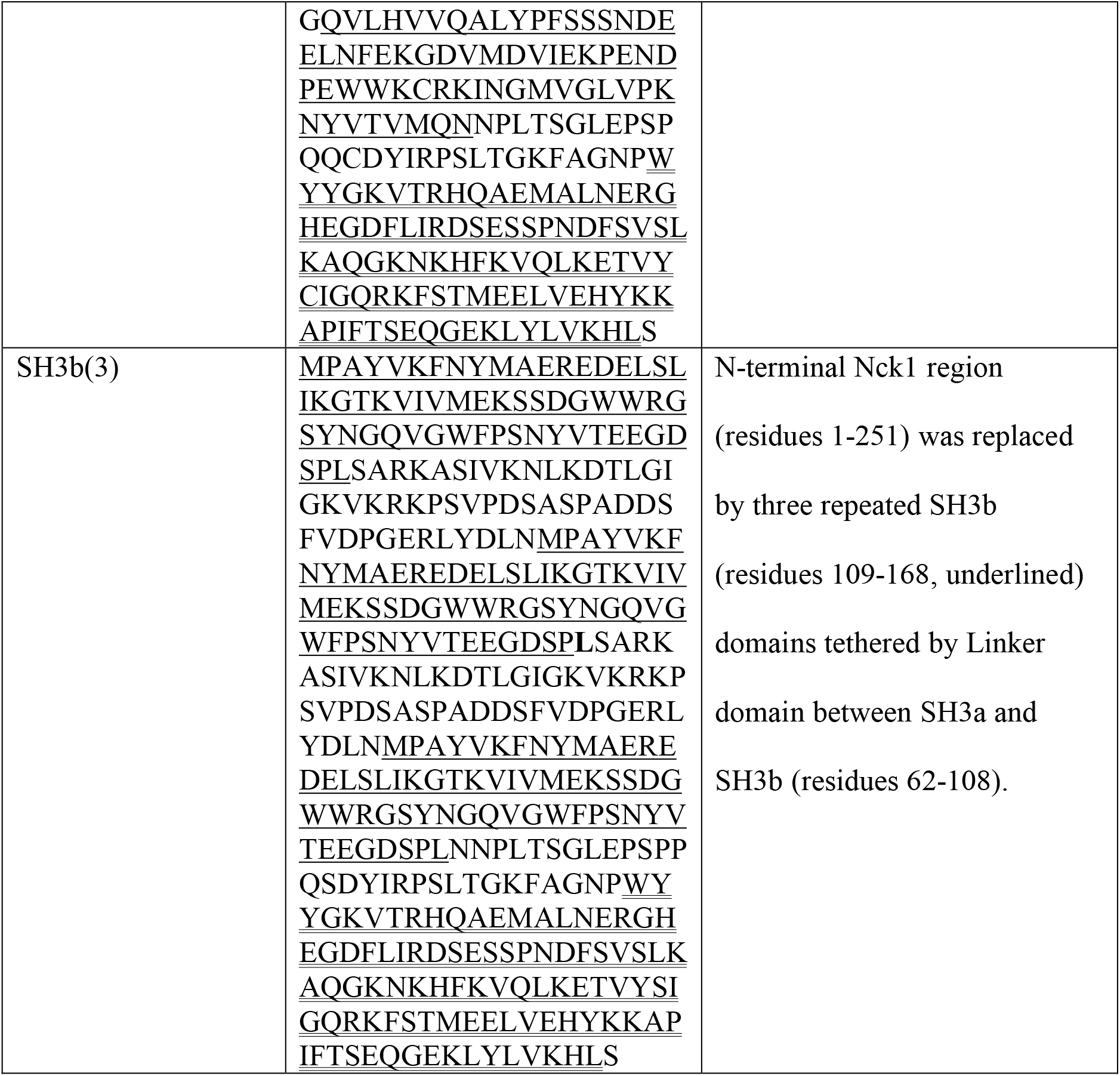
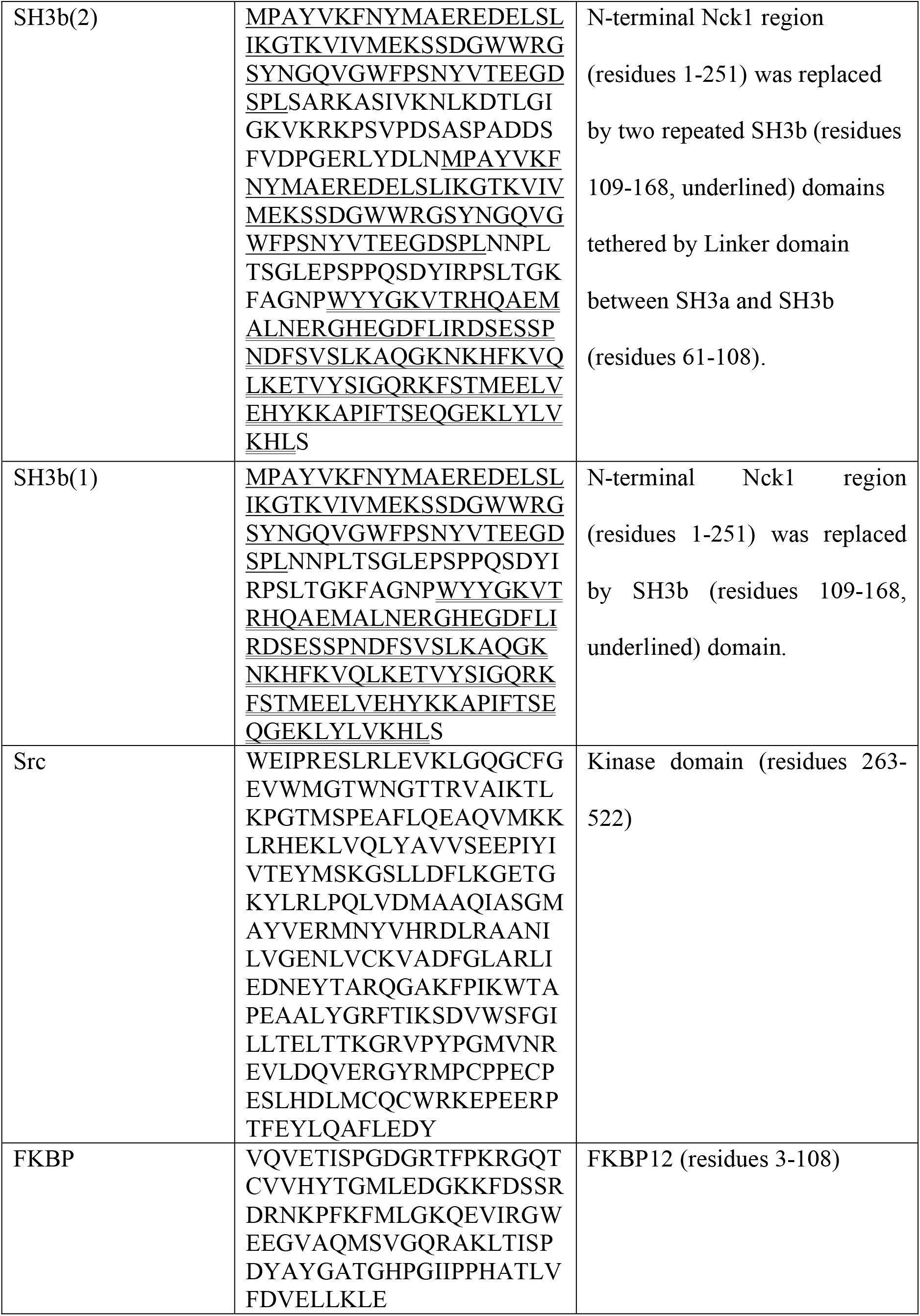
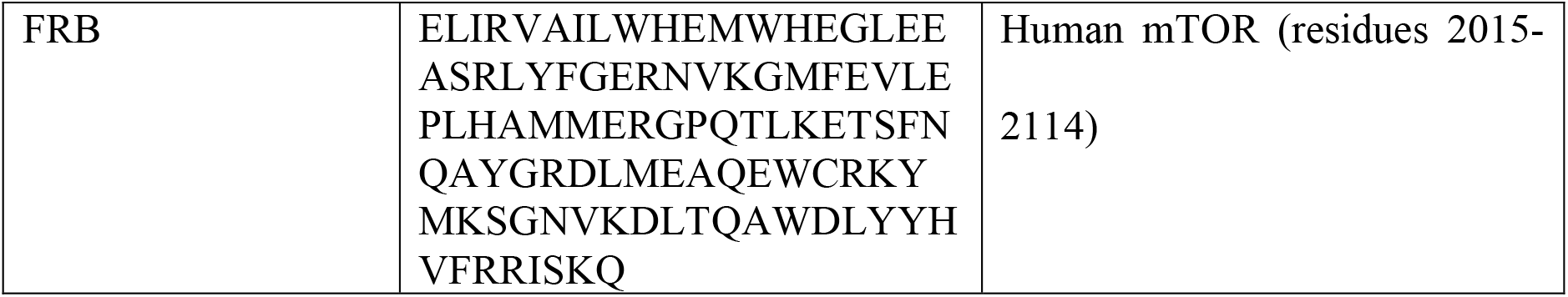
Information on protein expression constructs used in this study

### Cell Culture and transfection

#### Cell Culture

The HeLa cell line was kindly provided by the lab of Hongtao Yu (UT Southwestern Medical Center) and the HEK293 cell line was purchased from the ATCC (CRL-1573). Both cell lines were maintained in Dulbecco’s modified Eagle’s medium (Invitrogen) supplemented by 10% FBS (Hyclone), 1 mM penicillin/streptomycin (Invitrogen) and 1 mM GlutaMAX (Invitrogen) at 37°C with 5% CO_2_.

#### Transfection

Cells were transferred into 6-well plates (Fisher Scientific) for immunoblotting experiments or 35 mm glass-bottom dish (Mattek) for imaging experiments one day before transient transfection. Cell confluency at the point of transfection was 60∼80%. For TIRF imaging, to enhance even attachment of cells to the surface, 35 mm glass-bottom dishes (Mattek) were coated with poly-lysine (by incubating with 0.1% L-lysine (Sigma) solution for 30 min at room temperature followed by two washes with growth media) before cells were applied.

Effectene (QIAGEN) was used for transient transfection according to the manufacturer’s protocol with 0.4 ug of DNA for each sample. For co-expression of Src-FKBP, Nephrin-FRB, and Nck1 (or engineered Nck1 proteins), 0.1 ug, 0.2 ug, and 0.1 ug of DNA were used, respectively. When either cSrc-FKBP or Nck1 was not expressed, empty pcDNA vector (Invitrogen) was used to match the total DNA amount. Samples were incubated for 27 h at 37 °C prior to experiments to ensure sufficient expression and membrane localization of Nephrin-FRB. Experiments were done using cells incubated no longer than 35 h after transfection to avoid cell damage from unregulated Src kinase activity and overexpressed Nck1.

### Coimmunoprecipitation and Western Blotting

HeLa and HEK293 cell samples prepared in 6-well plates (Fisher Scientific) were used for the assay. When required, rapamycin (Toronto Research, final concentration 1 uM) or a matching volume of DMSO was added to the media for 15 min as indicated in the figure legends. After removing the culture medium and washing cells two times with PBS, cells were harvested with 1 mM EDTA in PBS by scraping and pipetting. Cell suspensions were centrifuged at 10,000 g for 2 min at 4 °C to collect the cell pellets. Cell lysates were made by resuspending cell pellets with lysis buffer (50 mM Tris, pH 8.0, 150 mM NaCl, 1 % TritonX-100, 1 mM PMSF, 1 mM benzamidine, 1 µg/ml leupeptin, 1 µg/ml antipain) for 30 min on ice. To sustain the phosphorylation level of Nephrin-FRB, 10 mM NaF, 5 mM beta glycerophosphate, 2.5 mM Na_3_VO_4_ and 5 µM Okadaic acid were added to the buffer during lysis to inhibit phosphatases. After centrifugation at 16,000 g for 15 min at 4 °C, supernatants were collected in new tubes and immunoprecipitated with anti-Flag M2 affinity gel (Sigma) for 2 h at 4 °C. Immune complexes were washed two times with lysis buffer and eluted with 0.5 mg/mL Flag peptide (Sigma) in lysis buffer. Eluted samples were mixed with SDS sample buffer and heated at 95 °C for 10 min for SDS-PAGE and western blotting. Antibodies used in western blotting were anti-phosphotyrosine (4G10) (Millipore, 1:1000), anti-Nck (BD Biosciences, 1:1000), anti-Flag (Sigma, 1:2000) to identify Nephrin), anti-HA (Cell Signaling, 1:1000). For western blotting with 4G10 antibody, membranes were blocked and blotted with 5% BSA (Sigma) in Tris-buffered saline (TBS). Membranes were incubated with the primary antibodies overnight at 4 °C before washing and staining with secondary antibody.

### Cell preparation for imaging

#### Rapamycin-mediated Nephrin phosphorylation and chemical treatment

Transiently transfected cells in 35 mm glass-bottom dishes were starved with FBS-free media (Dulbecco’s modified Eagle’s medium (Invitrogen) supplemented by 1 mM penicillin/streptomycin (Invitrogen) and 1 mM GlutaMAX (Invitrogen)) for 7 h prior to imaging. At 27 h after transient transfection, methyl-beta-cyclodextrin (MβCD, Sigma, 1 mg/mL) was added to the culture media. After 10 min, cells were treated with rapamycin (Toronto Research, 2 µM) for 30 min. For the experiments with actin cytoskeleton perturbation, Latrunculin B (Sigma, 5 µM) was added to the culture media for 10 min before 20 min of rapamycin. For cluster motility assays, applying drugs indicated in transfected cells were transferred into Hank’s balanced salt solution (HBSS) supplemented with 0.5% BSA prior to chemical treatments. Cells were then treated for 20 min with Latrunculin B, blebbistatin S or R (50 µM, Toronto Research), CK666 or CK869 (100 µM, Millipore) after 20 min of rapamycin treatment. For peripheral cluster formation assays, cells were treated for 10 min of MβCD followed by 10 min of rapamycin.

#### Cell fixation and staining

For fixed cell experiments, after the treatments above cells were washed twice with Hank’s balanced salt solution (HBSS) and incubated with 4% paraformaldehyde (Sigma) in phosphate buffered saline (PBS) for 20 min at room temperature followed by two washes with PBS.

For immunofluorescence experiments, fixed cell samples were permeabilized with 0.1% saponin (Sigma) in PBS for 5 min. The antibodies used for staining were anti-clathrin heavy chain (provided by D.Billadeau and T.Gomez, 1:1000), anti-caveolin-1 (BD Transduction, 1:500), and anti-paxillin (Abcam, 1:1000) antibodies. Secondary antibodies conjugated with Alexa Fluor 647 (Invitrogen) were used. For F-actin staining, phalloidin conjugated with Alexa 647 (Invitrogen) was used.

### Epifluorescence microscopy imaging

Epifluorescence images of live cells (Figure 1B; Supplemental Figure S1A) or fixed cells (Supplemental Figures S1D and S6A) were acquired on an Applied Precision Deltavision RT microscope equipped with CoolSnap-HQ (Photometrics) CCD camera with a 60 X objective. In Figure 1B, line scanning profiles were measured using ImageJ (FIJI).

### TIRF microscopy imaging and data analysis

#### Nephrin/Nck cluster formation at the basal membrane in response to rapamycin

TIRF images for Figure 2A were acquired from live cells using a Leica DMI6000 microscope equipped with a Hamamatsu ImagEM X2 EM-CCD camera, a 100 x 1.49 NA objective and TIRF/iLAS2 module. All other TIRF images for Nephrin cluster formation were collected on a Nikon Eclipse Ti microscope equipped with an Andor iXon Ultra 897 EM-CCD camera with a 100 X objective in TIRF- or Epi-fluorescence mode. Live cells were imaged for Supplemental Figure S3A, while Figures 2, B-E, Supplemental Figure S2, Figure 3, Supplemental Figures S3, C-G, Figures 4, B and C, Supplemental Figure S4 were from fixed cell samples. All the images for the quantitative analysis of Nephrin cluster formation were from fixed samples and acquired with the same microscopic parameters (TIRF angle, camera gain, laser power, exposure time for each channel).

To identify cells containing clusters that were enriched with both Nephrin and Nck1, the following procedures were performed using automated macros in ImageJ. First, images were processed with ImageJ (FIJI) to subtract background intensities for each pixel and to correct uneven illumination with an image of homogeneous fluorescent solution (mEGFP or mCherry in HEPES buffer). Second, ratio images were generated by dividing Nck1 images by Nephrin images. Third, Nephrin channel (mCherry) images were segmented using Otsu thresholding in ImageJ to identify cell regions that evenly attached to the surface. Fourth, a representative cell area (RCA) was selected within each cell using the Maximum Inscribed Circles Plugin in ImageJ (developed by Olivier Burri from École Polytechnique Fédérale de Lausanne). Fifth, RenyiEntropy thresholding in ImageJ was applied to the RCAs and result areas larger than 4 pixels were classified as the regions of interest (ROIs). Sixth, intensities and sizes of Nephrin (mCherry channel in TIRF mode), Nck1 at the membrane (mEGFP channel in TIRF mode), Nck1 in the cytoplasm (mEGFP channel in Epifluorescence mode), and Nck1/Nephrin ratio (ratio images, mCherry channel divided by mEGFP channel, both in TIRF modes) were measured for the RCAs and for the ROIs. For each cell we then calculated the ratio of the mean Nephrin intensities from the ROIs to the mean of the sub-regions within RCAs excluding ROIs (Ratio#1), the ratio of the size of ROIs to the size of the RCA (Ratio#2), the mean of Nck1/Nephrin ratios within the ROIs (Ratio#3).

In order to train the automated scoring macro, we first manually scored a test group of cells from the Rap(WT) and Rap(Y3F) conditions (50 randomly selected cells each from 3 independent transfections) for the presence or absence of clusters. This manual scoring was used to determine cutoff values for Ratio#1 (≥ 1.45) and Ratio#2 (≥ 0.04; < 0.7) that minimized the sum of false positive/negative decisions by the macro. Additionally, we reasoned that Nephrin clusters forming as a result of Nck binding should be enriched in both proteins. For cells expressing Nephrin(Y3F), where Nephrin is not appreciably phosphorylated and thus does not bind Nck1, 90% of cells in the test set had ROIs with mean Nck1:Nephrin ratio (Ratio#3) < 0.9, indicating that this value corresponds to Nck1-independent clustering of Nephrin. Supplemental Figure S2C shows the fraction of cells with Nck1-dependent Nephrin clusters from those with Nephrin clusters, calculated by dividing the number of cells with Ratio#1 ≥ 1.45, 0.04 ≤ Ratio#2 < 0.7, Ratio #3 ≥ 0.9 by the number of cells with Ratio#1 ≥ 1.45, 0.04 ≤ Ratio#2 < 0.7 from the entire dataset (> 440 cells each condition) for each control condition (cells expressing Nephrin (Y3F), cells expression Nephrin (Y5F), cells without rapamycin treatment, cells without Src-FKBP expression). Thus, we classified Nck1-dependent clusters as those with Ratio#1 (≥ 1.45), Ratio#2 (≥ 0.04; < 0.7), and Ratio#3 (≥ 0.9). To calculate the fraction of cells with Nck1-dependent Nephrin clusters, we divided the number of cells containing Nck1-dependent clusters by the total number of cells analyzed.

We used logistic regression tests in MatLab to fit the probability of cluster formation for individual cells as a function of Nephrin-FRB or Nck1 expression level. Cell data in each condition (cell classifications as containing clusters or not; average protein intensities across the RCA from TIRF images for Nephrin and epi-fluorescence images for Nck1) were fitted to a single population distribution over either Nephrin-FRB or Nck1 protein intensities. To determine whether two samples were significantly different, a Chi-squared test was performed to report the p-value to reject the null hypothesis that the combined data set of two samples fit to a single-population distribution, while the alternative hypothesis is to fit to a two-population distribution.

#### Movement of protein clusters (Figure 5 and Supplemental Figure S5)

TIRF images for Nephrin cluster movement were acquired using a Leica DMI6000 microscope equipped with a Hamamatsu ImagEM X2 EM-CCD camera, a 100 x 1.49 NA objective and TIRF/iLAS2 module. All time-lapse images were collected with cells at 37 °C under 5% CO_2_ atmosphere. Images were collected every 250 msec for 30 sec. Microscope settings were optimized to minimize photo-damage to cells.

Cell boundaries were initially detected by applying Otsu thresholding in ImageJ. The diameter of the detected region was then reduced by 20 pixels to avoid tracking artifacts at of the cell periphery. These pre-processed time-lapse images were then analyzed using the ‘Track surface (over time)’ method from Imaris (Bitplane). Protein clusters were detected by thresholding with background subtraction and local contrast provided by Imaris. Any clusters smaller than 0.5 μm^2^ or larger than 100 μm^2^ were excluded from analysis to avoid either artifactual tracking of noise or high-speed data from centroid shifting of large sized clusters, respectively. Tracking analysis was done with a Brownian motion algorithm with the maximum moving distance of 0.5 μm between two consecutive frames (250 ms) for clusters that had duration times longer than 10 sec. The maximum gap size was set to 1 frame. The tracking data were processed to calculate mean-square-displacement (MSD) values using the Xtension ‘Mean-square-displacement analysis’ (Tarantino, Tinevez et al., 2014). The MSD values for the first ¼ of the track timecourse were used to calculate the diffusion coefficients and alpha value using Prism (Graphpad).

### STORM microscopy (Figure 2B)

Cells were transfected with Nephrin-FRB conjugated with mEOS2 along with Src-FKBP and Nck1 proteins. STORM images were acquired on fixed cell samples on a Nikon Eclipse Ti microscope equipped with an Andor iXon Ultra 897 EM-CCD camera and a 100 X objective in TIRF mode (Edelstein, Tsuchida et al., 2014). Images were processed with the Localization Microscopy plugin of Micromanager. More than 200,000 images were used to generate a STORM image followed by correction for X-Y drifting. The resulting image was rendered with points that are 25% of the size of raw image pixels.

### Fluorescence Recovery After Photobleaching (FRAP) (Figures 4D and 6D; Supplemental Figure S6B)

FRAP images for the clusters at the basal membrane or the cell periphery were imaged on live cells using a Zeiss LSM510 confocal microscope equipped with an EM-CCD camera and a 100X or 63X objective, respectively. For clusters at the basal membrane, a circle with 1.75 μm diameter was bleached for clustered and unclustered areas within the same cell, and recoveries were imaged every 3 sec over 180 sec. For clusters at the cell periphery, square areas of 2.23 μm x 2.23 μm were bleached and recoveries were imaged every 1 sec over 120 sec. Recovery data were corrected for background photobleaching using the time-lapse images of an unbleached cell and normalized with the pre-bleaching images, respectively (Dundr & Misteli, 2003, McNally, 2008). The mean values of normalized data were fit to either a single or a double-exponential decay function, as determined using an F-test implemented in Prism (Graphpad).

### Homo-FRET experiment (Figures 4, G and H)

Cell images for steady-state homo-FRET-based anisotropy measurements were acquired on live cells expressing Src-FKBP, Nephrin-FRB-mEYFP, and mCherry-Nck1. Images were acquired on a Nikon Eclipse Ti microscope equipped with an Andor TuCam dual camera arrangement and a 100 X objective in TIRF mode. Cells were excited with a polarization-preserving 488 nm laser, from which fluorescence emission was resolved into parallel and perpendicular polarized light through a polarizing beam splitter (Moxtek Inc, USA), and recorded with the two Andor iXon Ultra 897 EM-CCD cameras.

G-factor images were calculated by ImageJ using images of fluorescein solution (Goswami et al., 2008). G-factor and cell images were background corrected with images of water. MATLAB (MathWorks) was used to align the images pixel-by-pixel from the two cameras based on images of sub-resolution fluorescent beads, to correct perpendicular images using the G-factor images, and to obtain anisotropy maps smoothed by a spatial averaging filter (Ghosh et al, 2012; Saha et al 2015). The intensity and anisotropy levels were measured over time from 3.2 μm x 3.2 μm square regions either within areas containing protein clusters or in areas lacking clusters. MATLAB (MathWorks) was used to calculate the fraction of photobleaching at each time point and to bin the data for plotting.

### Spinning Disk confocal microscope imaging and data analysis (Figures 6, A-C)

Fixed cell images were acquired using a Perkin Elmer Spinning Disk Ultraview ERS equipped with a Hamamatsu C9100-50 EM-CCD camera to analyze Nephrin clustering at the cell periphery.

The following procedures were performed by an automated macro in ImageJ (FIJI) to identify cells with protein clusters at the periphery that were enriched with Nephrin, Nck and F-actin. First, background intensities for each channel were subtracted from cell images. Second, cell boundaries were detected using Mean thresholding in ImageJ on Nephrin channel images. Third, nucleus regions were masked using DAPI channel images due to the partial nuclear localization of the engineered Nck1 constructs. Fourth, using the Membrane Profile Plugin in ImageJ, the cells were divided into five degree radial sections and the peripheral membrane regions (20 pixel thickness) of each were determined. The mean intensities of Nephrin, Nck1 and F-actin within the membrane region of each radial section were measured (Sinha, Gao et al., 2010). Fifth, the radial intensities of Nephrin were segmented using Otsu thresholding in MatLab and the ratio of the average radial intensities above and below the thresholding cutoff was calculated (P-Ratio).

In order to train the automated cluster identification macro, we first manually scored a test group of cells from Rap(WT) and Rap(Y3F) transfections (the first set of cells from 3 independent transfections for each protein) for the presence or absence of clusters. This manual scoring was used to determine a cutoff value of P-Ratio ≥ 1.7, which resulted in the minimum number of false positive/negative decisions by the automated macro. To determine the enrichment of Nck1 and F-actin within the clusters, the correlation values of the radial intensities of Nephrin:Nck1 (COR#1) and Nephrin:F-actin (COR#2) were calculated, respectively. For cells expressing Nephrin(Y3F), where Nck1 does not bind to Nephrin, 80% of the cells in the test set had COR#1 < 0.7. Additionally, in the test set of Rap(WT), where interactions between Nephrin and Nck1 result in actin polymerization, more than 90% of cells that were manually scored positive had COR#2 ≥ 0.65. Thus, in automated cluster identification on the complete datasets in all conditions, we classified cells containing protein clusters at the cell periphery as those with P-Ratio ≥ 1.7, COR#1 ≥ 0.7, and COR#2 ≥ 0.65. In Figure 6C, cells expressing similar level of engineered Nck1 constructs (S3, S2 or S1) were analyzed.

## Acknowledgements

We thank Hongtao Yu (UTSW) for providing the HeLa cell line used in this work, Dan Billadeau and Timothy Gomez (Mayo Clinic) for providing antibodies, Nico Stuurman (UCSF) for assistance with STORM imaging, Kate Luby-Phelps and Abhijit Bugde (UTSW Live cell Imaging Core Facility) for their assistance in epi-fluorescence and spinning disk confocal experiments, Sudeep Banjade for advice on designing the S3, S2, S1 constructs, Khuloud Jaqaman (UTSW) for advice on cluster motility analysis, Salman Banani and Jonathan Ditlev (UTSW) for critical reading of the manuscript, and members of the Rosen Lab and participants in the MBL/HHMI Summer Institutes for advice and helpful discussions. This work was supported by a Howard Hughes Medical Institute Collaborative Innovation Award, the Welch Foundation (I-1544 to M.K.R.), a J.C. Bose Fellowship from the Department of Science and Technology, government of India (to S.M.), a Margadarshi Fellowship from the Wellcome Trust – Department of Biotechnology, India Alliance (IA/M/15/1/502018 to S.M.). Research in the Rosen lab is supported by the Howard Hughes Medical Institute.

## References

Altschuler SJ, Angenent SB, Wang Y, Wu LF (2008) On the spontaneous emergence of cell polarity. Nature 454: 886–9

Astro V, de Curtis I (2015) Plasma membrane-associated platforms: dynamic scaffolds that organize membrane-associated events. Sci Signal 8: re1

Banani SF, Rice AM, Peeples WB, Lin Y, Jain S, Parker R, Rosen MK (2016) Compositional Control of Phase-Separated Cellular Bodies. Cell 166: 651–663

Banaszynski LA, Liu CW, Wandless TJ (2005) Characterization of the FKBP.rapamycin.FRB ternary complex. J Am Chem Soc 127: 4715–21

Banjade S, Rosen MK (2014) Phase transitions of multivalent proteins can promote clustering of membrane receptors. Elife 3

Banjade S, Wu Q, Mittal A, Peeples WB, Pappu RV, Rosen MK (2015) Conserved interdomain linker promotes phase separation of the multivalent adaptor protein Nck. Proc Natl Acad Sci U S A 112: E6426–35

Barrantes FJ, Bermudez V, Borroni MV, Antollini SS, Pediconi MF, Baier JC, Bonini I, Gallegos C, Roccamo AM, Valles AS, Ayala V, Kamerbeek C (2010) Boundary lipids in the nicotinic acetylcholine receptor microenvironment. J Mol Neurosci 40: 87–90

Blasutig IM, New LA, Thanabalasuriar A, Dayarathna TK, Goudreault M, Quaggin SE, Li SS, Gruenheid S, Jones N, Pawson T (2008) Phosphorylated YDXV motifs and Nck SH2/SH3 adaptors act cooperatively to induce actin reorganization. Mol Cell Biol 28: 2035–46

Buday L, Wunderlich L, Tamas P (2002) The Nck family of adapter proteins: regulators of actin cytoskeleton. Cell Signal 14: 723–31

Day CA, Kenworthy AK (2009) Tracking microdomain dynamics in cell membranes. Biochim Biophys Acta 1788: 245–53

Ditlev JA, Vega AR, Köster DV, Su X, Lakoduk A, Vale RD, Mayor S, Jaqaman K, Rosen MK (2018) A Composition-Dependent Molecular Clutch Between T Cell Signaling Clusters and Actin. bioRxiv

Dundr M, Misteli T (2003) Measuring dynamics of nuclear proteins by photobleaching. Curr Protoc Cell Biol Chapter 13: Unit 13 5

Edelstein AD, Tsuchida MA, Amodaj N, Pinkard H, Vale RD, Stuurman N (2014) Advanced methods of microscope control using muManager software. J Biol Methods 1

Edidin M (2003) Lipids on the frontier: a century of cell-membrane bilayers. Nat Rev Mol Cell Biol 4: 414–8

Flory PJ (1953) Principles of polymer chemistry. Cornell University Press, Ithaca,

Foda ZH, Shan Y, Kim ET, Shaw DE, Seeliger MA (2015) A dynamically coupled allosteric network underlies binding cooperativity in Src kinase. Nat Commun 6: 5939

Fromm SA, Kamenz J, Noldeke ER, Neu A, Zocher G, Sprangers R (2014) In vitro reconstitution of a cellular phase-transition process that involves the mRNA decapping machinery. Angew Chem Int Ed Engl 53: 7354–9

Gerke P (2003) Homodimerization and Heterodimerization of the Glomerular Podocyte Proteins Nephrin and NEPH1. Journal of the American Society of Nephrology 14: 918–926

Ghosh S, Saha S, Goswami D, Bilgrami S, Mayor S (2012) Dynamic imaging of homo-FRET in live cells by fluorescence anisotropy microscopy. Methods Enzymol 505: 291–327

Goswami D, Gowrishankar K, Bilgrami S, Ghosh S, Raghupathy R, Chadda R, Vishwakarma R, Rao M, Mayor S (2008) Nanoclusters of GPI-anchored proteins are formed by cortical actin-driven activity. Cell 135: 1085–97

Gowrishankar K, Ghosh S, Saha S C R, Mayor S, Rao M (2012) Active remodeling of cortical actin regulates spatiotemporal organization of cell surface molecules. Cell 149: 1353–67

Grahammer F, Wigge C, Schell C, Kretz O, Patrakka J, Schneider S, Klose M, Arnold SJ, Habermann A, Brauniger R, Rinschen MM, Volker L, Bregenzer A, Rubbenstroth D, Boerries M, Kerjaschki D, Miner JH, Walz G, Benzing T, Fornoni A et al. (2016) A flexible, multilayered protein scaffold maintains the slit in between glomerular podocytes. JCI Insight 1

Grecco HE, Schmick M, Bastiaens PI (2011) Signaling from the living plasma membrane. Cell 144: 897–909

Greka A, Mundel P (2012) Cell biology and pathology of podocytes. Annu Rev Physiol 74: 299–323

Groves JT, Kuriyan J (2010) Molecular mechanisms in signal transduction at the membrane. Nat Struct Mol Biol 17: 659–65

Honerkamp-Smith AR, Veatch SL, Keller SL (2009) An introduction to critical points for biophysicists; observations of compositional heterogeneity in lipid membranes. Biochim Biophys Acta 1788: 53–63

Hong J, Murugesan S, Betzig E, Hammer JA (2017) Contractile actomyosin arcs promote the activation of primary mouse T cells in a ligand-dependent manner. PLoS One 12: e0183174

Jaqaman K, Grinstein S (2012) Regulation from within: the cytoskeleton in transmembrane signaling. Trends Cell Biol 22: 515–26

Jones N, Blasutig IM, Eremina V, Ruston JM, Bladt F, Li H, Huang H, Larose L, Li SS, Takano T, Quaggin SE, Pawson T (2006) Nck adaptor proteins link nephrin to the actin cytoskeleton of kidney podocytes. Nature 440: 818–23

Jones N, New LA, Fortino MA, Eremina V, Ruston J, Blasutig IM, Aoudjit L, Zou Y, Liu X, Yu GL, Takano T, Quaggin SE, Pawson T (2009) Nck proteins maintain the adult glomerular filtration barrier. J Am Soc Nephrol 20: 1533–43

Kestila M, Lenkkeri U, Mannikko M, Lamerdin J, McCready P, Putaala H, Ruotsalainen V, Morita T, Nissinen M, Herva R, Kashtan CE, Peltonen L, Holmberg C, Olsen A, Tryggvason K (1998) Positionally cloned gene for a novel glomerular protein--nephrin--is mutated in congenital nephrotic syndrome. Mol Cell 1: 575–82

Kholodenko BN, Hoek JB, Westerhoff HV (2000) Why cytoplasmic signalling proteins should be recruited to cell membranes. Trends Cell Biol 10: 173–8

Khoshnoodi J, Sigmundsson K, Ofverstedt LG, Skoglund U, Obrink B, Wartiovaara J, Tryggvason K (2003) Nephrin promotes cell-cell adhesion through homophilic interactions. Am J Pathol 163: 2337–46

Koster DV, Husain K, Iljazi E, Bhat A, Bieling P, Mullins RD, Rao M, Mayor S (2016) Actomyosin dynamics drive local membrane component organization in an in vitro active composite layer. Proc Natl Acad Sci U S A 113: E1645–54

Koster DV, Mayor S (2016) Cortical actin and the plasma membrane: inextricably intertwined. Curr Opin Cell Biol 38: 81–9

Kusumi A, Suzuki KG, Kasai RS, Ritchie K, Fujiwara TK (2011) Hierarchical mesoscale domain organization of the plasma membrane. Trends Biochem Sci 36: 604–15

Lenkkeri U, Mannikko M, McCready P, Lamerdin J, Gribouval O, Niaudet PM, Antignac CK, Kashtan CE, Homberg C, Olsen A, Kestila M, Tryggvason K (1999) Structure of the gene for congenital nephrotic syndrome of the finnish type (NPHS1) and characterization of mutations. Am J Hum Genet 64: 51–61

Li H, Zhu J, Aoudjit L, Latreille M, Kawachi H, Larose L, Takano T (2006) Rat nephrin modulates cell morphology via the adaptor protein Nck. Biochem Biophys Res Commun 349: 310–6

Li P, Banjade S, Cheng HC, Kim S, Chen B, Guo L, Llaguno M, Hollingsworth JV, King DS, Banani SF, Russo PS, Jiang QX, Nixon BT, Rosen MK (2012) Phase transitions in the assembly of multivalent signalling proteins. Nature 483: 336–40

Lin Y, Protter DS, Rosen MK, Parker R (2015) Formation and Maturation of Phase-Separated Liquid Droplets by RNA-Binding Proteins. Mol Cell 60: 208–19

Martin CE, Jones N (2018) Nephrin Signaling in the Podocyte: An Updated View of Signal Regulation at the Slit Diaphragm and Beyond. Front Endocrinol (Lausanne) 9: 302

Martin GS (2001) The hunting of the Src. Nat Rev Mol Cell Biol 2: 467–75

McNally JG (2008) Quantitative FRAP in Analysis of Molecular Binding Dynamics In Vivo. 85: 329–351

Mitrea DM, Cika JA, Guy CS, Ban D, Banerjee PR, Stanley CB, Nourse A, Deniz AA, Kriwacki RW (2016) Nucleophosmin integrates within the nucleolus via multi-modal interactions with proteins displaying R-rich linear motifs and rRNA. Elife 5

Murugesan S, Hong J, Yi J, Li D, Beach JR, Shao L, Meinhardt J, Madison G, Wu X, Betzig E, Hammer JA (2016) Formin-generated actomyosin arcs propel T cell receptor microcluster movement at the immune synapse. J Cell Biol 215: 383–399

New LA, Keyvani Chahi A, Jones N (2013) Direct regulation of nephrin tyrosine phosphorylation by Nck adaptor proteins. J Biol Chem 288: 1500–10

New LA, Martin CE, Scott RP, Platt MJ, Keyvani Chahi A, Stringer CD, Lu P, Samborska B, Eremina V, Takano T, Simpson JA, Quaggin SE, Jones N (2016) Nephrin Tyrosine Phosphorylation Is Required to Stabilize and Restore Podocyte Foot Process Architecture. J Am Soc Nephrol 27: 2422–35

Nikolov DB, Xu K, Himanen JP (2013) Eph/ephrin recognition and the role of Eph/ephrin clusters in signaling initiation. Biochim Biophys Acta 1834: 2160–5

Padrick SB, Rosen MK (2010) Physical mechanisms of signal integration by WASP family proteins. Annual review of biochemistry 79: 707–35

Paluch E, Sykes C, Prost J, Bornens M (2006) Dynamic modes of the cortical actomyosin gel during cell locomotion and division. Trends Cell Biol 16: 5–10

Pollard TD, Cooper JA (2009) Actin, a central player in cell shape and movement. Science 326: 1208–12

Qin XS, Tsukaguchi H, Shono A, Yamamoto A, Kurihara H, Doi T (2009) Phosphorylation of nephrin triggers its internalization by raft-mediated endocytosis. J Am Soc Nephrol 20: 2534–45

Rao M, Mayor S (2014) Active organization of membrane constituents in living cells. Curr Opin Cell Biol 29: 126–32

Ruotsalainen V, Ljungberg P, Wartiovaara J, Lenkkeri U, Kestila M, Jalanko H, Holmberg C, Tryggvason K (1999) Nephrin is specifically located at the slit diaphragm of glomerular podocytes. Proc Natl Acad Sci U S A 96: 7962–7

Schleich K, Warnken U, Fricker N, Ozturk S, Richter P, Kammerer K, Schnolzer M, Krammer PH, Lavrik IN (2012) Stoichiometry of the CD95 death-inducing signaling complex: experimental and modeling evidence for a death effector domain chain model. Mol Cell 47: 306–19

Sherman E, Barr V, Manley S, Patterson G, Balagopalan L, Akpan I, Regan CK, Merrill RK, Sommers CL, Lippincott-Schwartz J, Samelson LE (2011) Functional nanoscale organization of signaling molecules downstream of the T cell antigen receptor. Immunity 35: 705–20

Sinha SK, Gao N, Guo Y, Yuan D (2010) Mechanism of induction of NK activation by 2B4 (CD244) via its cognate ligand. J Immunol 185: 5205–10

Su X, Ditlev JA, Hui E, Xing W, Banjade S, Okrut J, King DS, Taunton J, Rosen MK, Vale RD (2016) Phase separation of signaling molecules promotes T cell receptor signal transduction. Science 352: 595–9

Tarantino N, Tinevez JY, Crowell EF, Boisson B, Henriques R, Mhlanga M, Agou F, Israel A, Laplantine E (2014) TNF and IL-1 exhibit distinct ubiquitin requirements for inducing NEMO-IKK supramolecular structures. J Cell Biol 204: 231–45

van Zanten TS, Mayor S (2015) Current approaches to studying membrane organization. F1000Res 4

Verma R, Kovari I, Soofi A, Nihalani D, Patrie K, Holzman LB (2006) Nephrin ectodomain engagement results in Src kinase activation, nephrin phosphorylation, Nck recruitment, and actin polymerization. J Clin Invest 116: 1346–59

Verma R, Venkatareddy M, Kalinowski A, Patel SR, Garg P (2016) Integrin Ligation Results in Nephrin Tyrosine Phosphorylation In Vitro. PLoS One 11: e0148906

Verma R, Wharram B, Kovari I, Kunkel R, Nihalani D, Wary KK, Wiggins RC, Killen P, Holzman LB (2003) Fyn binds to and phosphorylates the kidney slit diaphragm component Nephrin. J Biol Chem 278: 20716–23

Wartiovaara J, Ofverstedt LG, Khoshnoodi J, Zhang J, Makela E, Sandin S, Ruotsalainen V, Cheng RH, Jalanko H, Skoglund U, Tryggvason K (2004) Nephrin strands contribute to a porous slit diaphragm scaffold as revealed by electron tomography. J Clin Invest 114: 1475–83

Weisswange I, Newsome TP, Schleich S, Way M (2009) The rate of N-WASP exchange limits the extent of ARP2/3-complex-dependent actin-based motility. Nature 458: 87–91

Welsh GI, Saleem MA (2010) Nephrin-signature molecule of the glomerular podocyte? J Pathol 220: 328–37

Wu H (2013) Higher-order assemblies in a new paradigm of signal transduction. Cell 153: 287–92

Wu L, Pan L, Zhang C, Zhang M (2012) Large protein assemblies formed by multivalent interactions between cadherin23 and harmonin suggest a stable anchorage structure at the tip link of stereocilia. J Biol Chem 287: 33460–71

Wu Y, Vendome J, Shapiro L, Ben-Shaul A, Honig B (2011) Transforming binding affinities from three dimensions to two with application to cadherin clustering. Nature 475: 510–3

Yap AS, Gomez GA, Parton RG (2015) Adherens Junctions Revisualized: Organizing Cadherins as Nanoassemblies. Dev Cell 35: 12–20

Yi J, Wu XS, Crites T, Hammer JA, 3rd (2012) Actin retrograde flow and actomyosin II arc contraction drive receptor cluster dynamics at the immunological synapse in Jurkat T cells. Mol Biol Cell 23: 834–52

Yu Y, Smoligovets AA, Groves JT (2013) Modulation of T cell signaling by the actin cytoskeleton. J Cell Sci 126: 1049–58

Zeng M, Shang Y, Araki Y, Guo T, Huganir RL, Zhang M (2016) Phase Transition in Postsynaptic Densities Underlies Formation of Synaptic Complexes and Synaptic Plasticity. Cell 166: 1163–1175 e12

