## Supplementary Figures for "Phosphorylation of Nephrin induces phase separated domains that move through actomyosin contraction"

Correspondence to Michael K. Rosen

This PDF file includes:

Captions for Movies S1 to S6

Supplemental Figure S1 to S6

Other Supplementary Materials for this manuscript include the following:

Movies S1 to S5

**Movie S1. Nephrin/Nck1 clusters form in response to rapamycin treatment.** TIRF time-lapse images of a live HeLa cell expressing Src-FKBP, Nephrin-FRB (shown in video) and Nck1. Cells were treated with 10 min of MβCD followed by rapamycin. Images were captured every 30 s after adding rapamycin to the media.

**Movie S2. Nephrin/Nck1 cluster formation is independent of actin polymerization.** TIRF time-lapse images of a live HeLa cell expressing Src-FKBP, Nephrin-FRB (shown in video) and Nck1. Cells were treated with 10 min of MβCD, 15 min of latrunculin B followed by rapamycin. Images were captured every 30 s after adding rapamycin to the media.

**Movie S3. Nephrin/Nck1 clusters at the basal membrane are mobile.** TIRF time-lapse images of a live HeLa cell (Figure 5A, first row) expressing Src-FKBP, Nephrin-FRB (shown in video) and Nck1. Cells were treated with 10 min of MβCD, 30 min of rapamycin. Images in the video were captured every 1 sec after rapamycin treatment.

**Movie S4. Nephrin/Nck1 clusters stop moving upon latrunculin B treatment.** TIRF time-lapse images of a live HeLa cell (Figure 5A, second row) expressing Src-FKBP, Nephrin-FRB (shown in video) and Nck1. Cells were treated with 10 min of MβCD, 20 min of rapamycin followed by 20 min of latrunculin B. Images in the video were captured every 1 sec at the end of the latrunculin B incubation.

**Movie S5. Nephrin/Nck1 clusters stop moving upon blebbistatin treatment.** TIRF time-lapse images of a live HeLa cell (Figure 5A, third row) expressing Src-FKBP, Nephrin-FRB (shown in video) and Nck1. Cells were treated with 10 min of MβCD, 20 min of rapamycin followed by 20 min of blebbistatin. Images in the video were captured every 1 sec at the end of the blebbistatin incubation.

**Movie S6. Movement of Nephrin/Nck1 clusters is decreased by inhibition of the Arp2/3 complex.** TIRF time-lapse images of a live HeLa cell expressing Src-FKBP, Nephrin-FRB (shown in video) and Nck1. Cells were treated with 10 min of MβCD, 20 min of rapamycin followed by 20 min of CK666. Images in the video were captured every 1 sec at the end of the CK666 incubation.

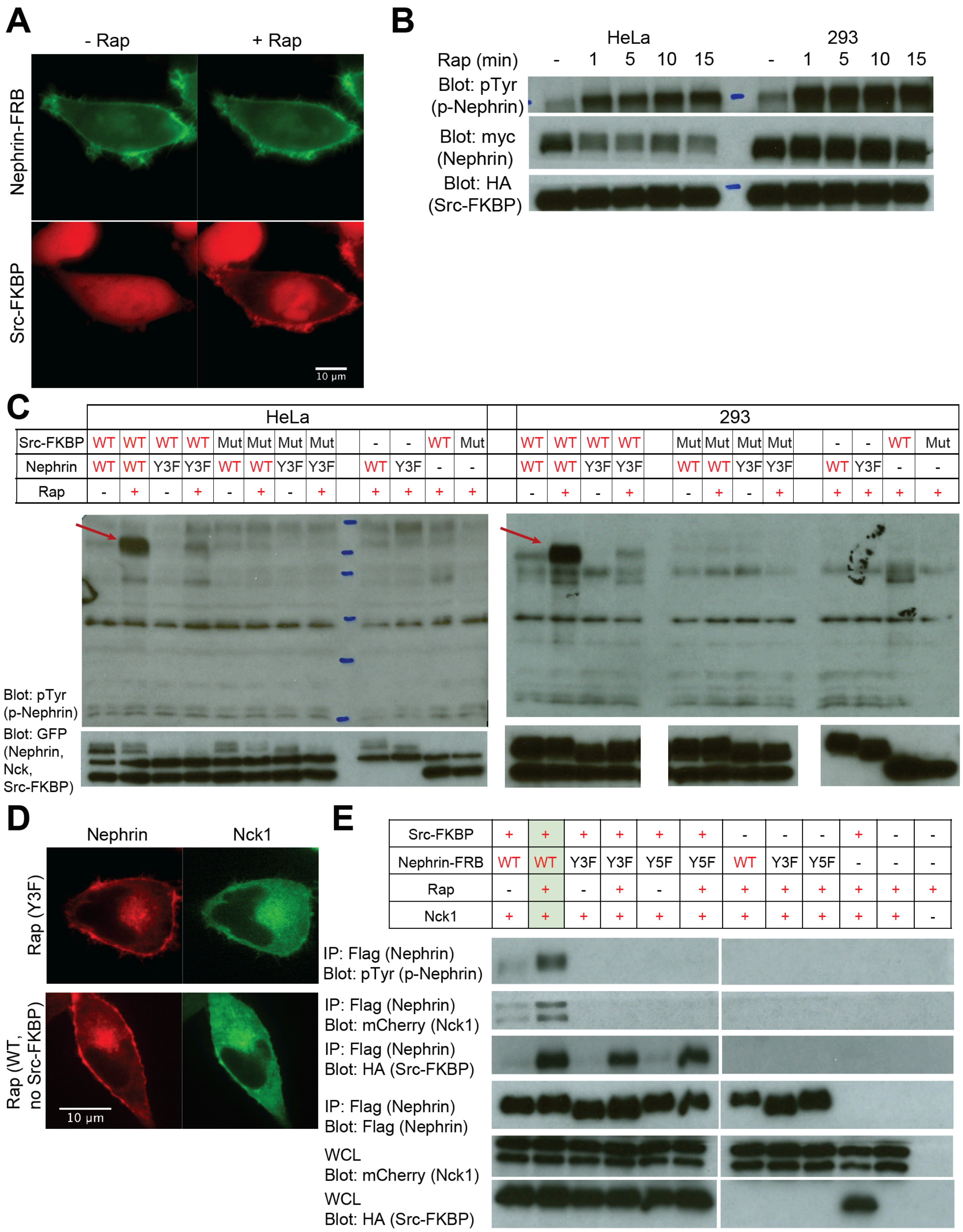

**Supplemental Figure S1. Rapamycin mediated Src binding leads to Nephrin phosphorylation.**

(A) Epi fluorescence images of a live HeLa cell expressing Src-FKBP (red), Nephrin-FRB (green) and Nck1 before (left column) or < 1 min after (right column) addition of rapamycin to the media.

(B) Western blots of HeLa (left five lanes) or 293 (right five lanes) whole cell lysates from cells expressing Src-FKBP, Nephrin-FRB, and Nck1. Antibodies were against pTyr (p-Nephrin), myc (Nephrin), or HA (Src-FKBP), as indicated. Cells were lysed after the indicated time of rapamycin treatment.

(C) Western blots of whole cell lysates from HeLa and 293 cells expressing a subset of Src(WT or inactive mutant (Mut) as indicated)-FKBP, Nephrin(WT, Y3F or Y5F as indicated)-FRB and Nck1. Antibodies were against pTyr (p-Nephrin, top) and EGFP (Nephrin, Nck1, and Src-FKBP) (bottom). Cells were treated with either DMSO (lanes 1, 3, 5, 7 in each gel) or rapamycin for 15 min before being lysed. Phosphorylated Nephrin-FRB is indicated with arrows based on the size of the construct.

(D) Spinning disk confocal images of fixed HeLa cells expressing Src-FKBP (top row only), Nephrin(WT or Y3F as indicated)-FRB (red) and Nck1 (green) 10 min after addition of rapamycin to the media. Images are shown with different contrast and brightness settings since Nck1 expression was much higher in most cells when Src-FKBP was not expressed.

(E) Western blots of 293 cells expressing a subset of Src-FKBP, Nephrin(WT, Y3F or Y5F as indicated)-FRB and Nck1. Antibodies were against pTyr (p-Nephrin), mCherry (Nck1), HA (Src-FKBP), Flag (Nephrin), as indicated. Either whole cell lysate (bottom two rows) or samples immunoprecipitated with anti-Flag affinity gel (top four rows) were used. Cells were treated with either DMSO (lanes 1, 3, 5) or rapamycin for 15 min before being lysed.

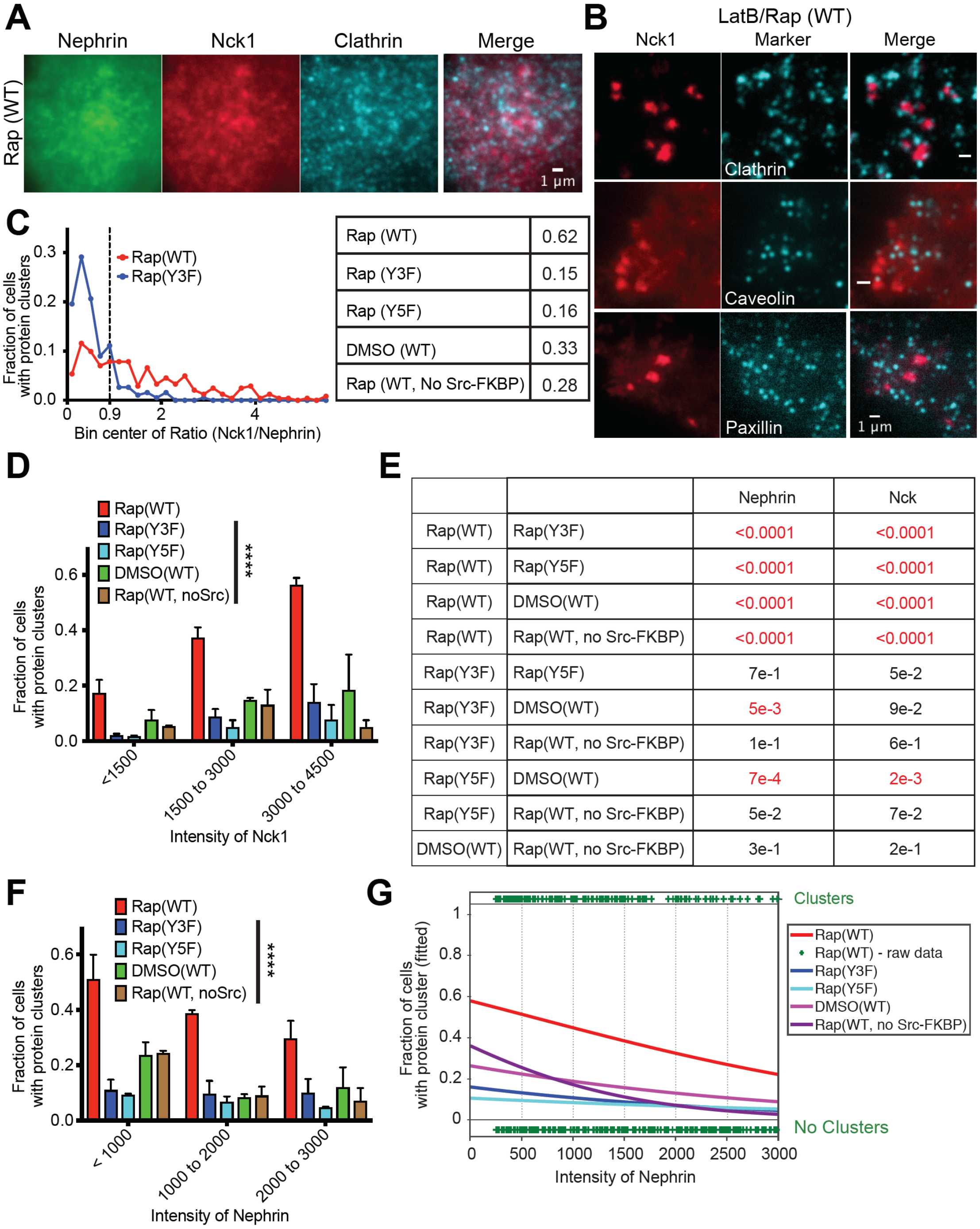

**Supplemental Figure S2. Nephrin/Nck1 clusters are not vesicular structures and form dependent on the cellular concentration of Nck1**

(A) TIRF images of a fixed HeLa cell expressing Src-FKBP, Nephrin-FRB (green), and Nck1 (red). Cells were immunofluorescence labeled to show clathrin (cyan). Right image shows a merge of the left three channels. Cells were treated with 10 min of MβCD followed by 30 min of rapamycin treatment prior to fixation.

(B) TIRF images of fixed HeLa cells expressing Src-FKBP, Nephrin-FRB, and Nck1. Left column shows Nck1. In middle column clathrin (top row), caveolin (middle row), or paxillin (bottom row) were detected by immunofluorescence. Right column shows a merged image of the left two channels. Cells were treated with 10 min of MβCD, 15 min of latrunculin followed by 15 min of rapamycin treatment prior to fixation.

(C to G) Quantitative analyses of protein clustering in various experimental conditions and at different cellular concentrations of Nck1. TIRF images of Nephrin and epi/TIRF images of Nck1 from fixed HeLa cells were used for the analysis. Cells expressed Src-FKBP (unless noted as 'no Src-FKBP'), Nck1, and Nephrin(WT, Y3F, or Y5F as indicated)-FRB. Cells were treated with 10 min of MβCD followed by either 30 min of rapamycin (Rap), 30 min of DMSO (DMSO) ($N = 3$ independent experiments, each with $> 140$ cell images).

(C) Plot of the fraction of cells with clusters versus the ratio of Nck1 and Nephrin fluorescence intensities. The table at right shows the fraction of cells containing Nephrin clusters with a ratio of Nck1/Nephrin TIRF intensities $> 0.9$ (indicated with reference line in the graph) in various experimental conditions.

(D and F) Graphs showing the fraction of cells with clusters for different expression levels of Nck1 (D) or Nephrin (F). Cell samples were binned according to the epi fluorescence intensity level of Nck1 (D) or the TIRF fluorescence intensity level of Nephrin (F). Error bars, s.e.m. (****$p < 0.0001$)

(E) Table reports the p-value calculated from a Chi-squared test when the null hypothesis is to fit a combined data set between two pairs into a single population distribution, while the alternative hypothesis is to fit into a two-population distribution. Experimental conditions with significant difference ($p < 0.01$) are colored red.

(G) Graphs showing the fitted probability of a cell forming clusters at a given Nephrin intensity level at the basal membrane. Cell data for each condition were fitted (lines) with a logistic regression model (left axis). Each plus sign indicates the status of a single cell (containing or lacking clusters) from 'Rap(WT)' sample (right axis).

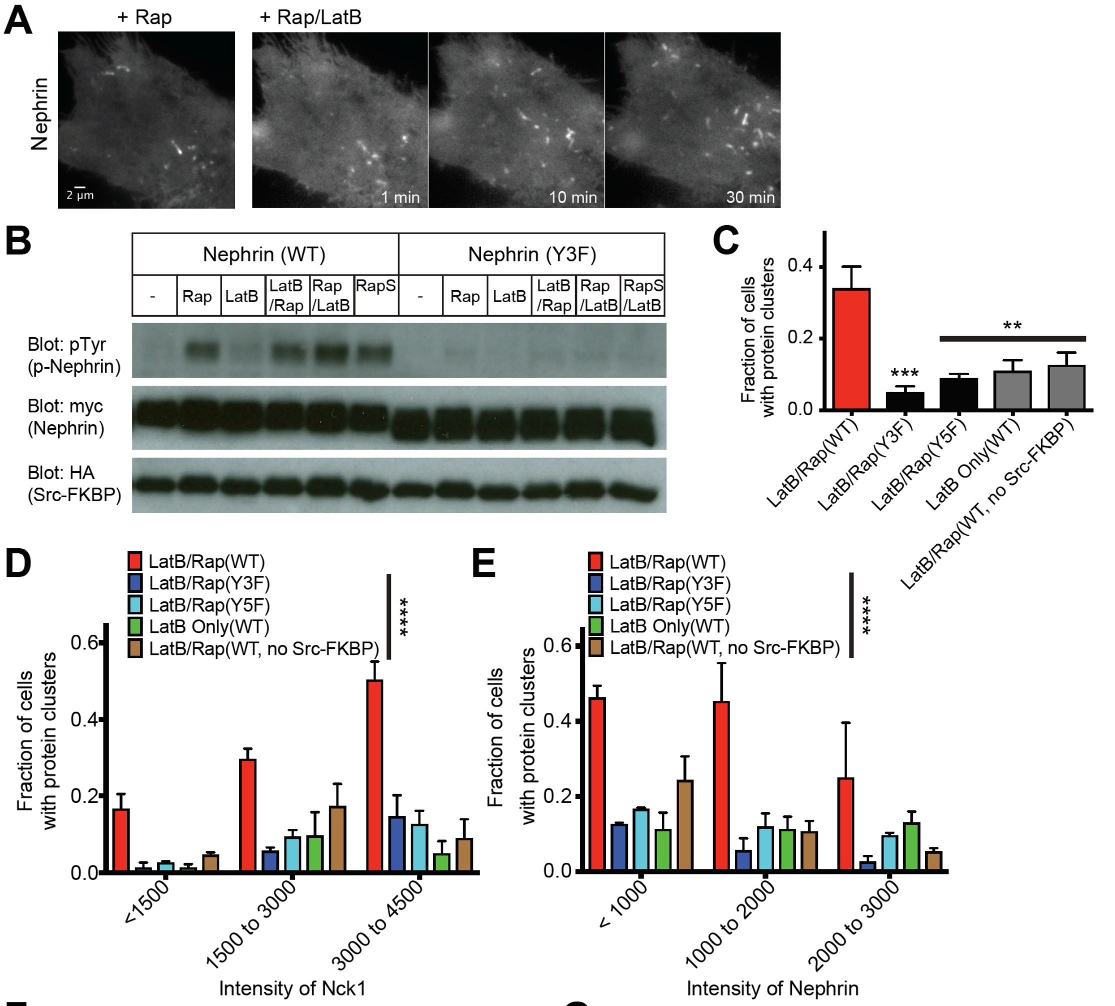

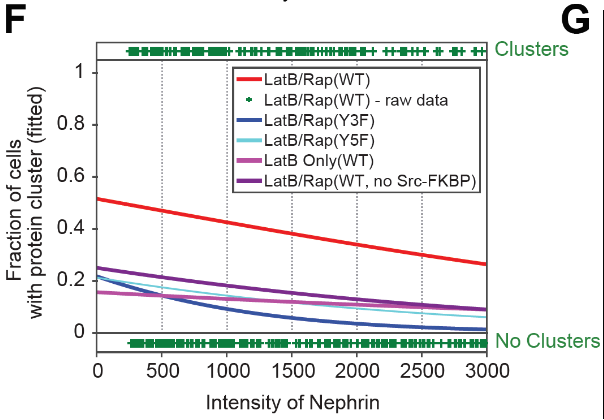

**G**

| | | Nephrin | Nck1 |
|---|---|---|---|
| LatB/Rap(WT) | LatB/Rap(Y3F) | <0.0001 | <0.0001 |
| LatB/Rap(WT) | LatB/Rap(Y5F) | <0.0001 | <0.0001 |
| LatB/Rap(WT) | LatB Only(WT) | <0.0001 | <0.0001 |
| LatB/Rap(WT) | LatB/Rap(WT, no Src-FKBP) | <0.0001 | <0.0001 |
| LatB/Rap(Y3F) | LatB/Rap(Y5F) | 1e-2 | 3e-1 |
| LatB/Rap(Y3F) | LatB Only(WT) | 3e-3 | 5e-1 |
| LatB/Rap(Y3F) | LatB/Rap(WT, no Src-FKBP) | <0.0001 | 4e-3 |
| LatB/Rap(Y5F) | LatB Only(WT) | 7e-1 | 8e-1 |
| LatB/Rap(Y5F) | LatB/Rap(WT, no Src-FKBP) | 1e-1 | 2e-1 |
| LatB Only(WT) | LatB/Rap(WT, no Src-FKBP) | 2e-1 | 3e-1 |

**Supplemental Figure S3. Nephrin/Nck1 cluster formation is independent of actin polymerization.**

(A) TIRF images of a live HeLa cell expressing Src-FKBP, Nephrin-FRB (shown in figure) and Nck1. The cell was treated for 10 min of MβCD and 15 min of rapamycin (left) followed by latrunculin B. Images taken at 1, 10, 30 min after latrunculin B treatment are shown in the right panel.

(B) Western blots of whole cell lysates from HeLa cells expressing Src-FKBP, Nephrin (WT or Y3F as indicated)-FRB and Nck1 against pTyr, myc, and HA as indicated. Cells were treated with either DMSO (-), 10 min of rapamycin (Rap), 10 min of latrunculin B (LatB), 10 min of latrunculin followed by 10 min of rapamycin (LatB/Rap), 10 min of rapamycin followed by 10 min of latrunculin (Rap/LatB), or 1 min of rapamycin followed by 10 min of latrunculin (RapS/LatB) prior to cell lysis.

(C to G) Quantitative analyses of protein clustering in various experimental conditions and at different cellular concentration of Nck1. TIRF images of Nephrin and epi/TIRF images of Nck1 from fixed HeLa cells were used for the analysis. Cells expressed Src-FKBP (unless noted as 'no Src-FKBP'), Nck1, and Nephrin(WT, Y3F, or Y5F as indicated)-FRB. Data noted as 'LatB Only' were from cells treated with 10 min of MβCD followed by 30 min of latrunculin B. All other data are from cells treated with 10 min of MβCD, 15 min of latrunculin B and 15 min of rapamycin prior to fixation. (N = 3 independent experiments, each with > 140 cell images).

(C to E) Error bars, s.e.m. (**$p < 0.01$, ***$p < 0.001$, ****$p < 0.0001$)

(C) Graph showing the fraction of cells with clusters in various experimental conditions.

(D and E) Graphs showing the fraction of cells with clusters expressing different levels of Nck1 (D) or Nephrin (E). Cell samples were binned according to the epi fluorescence intensity of Nck1 (C) or to the TIRF intensity of Nephrin (D).

(F) Graphs showing the fitted probability of a cell forming clusters at a given Nephrin intensity level at the basal membrane. Cell data for each condition were fitted (lines) with a logistic regression model (left axis). Each plus sign indicates the status of a single cell (containing or lacking clusters) from 'Rap(WT)' sample (right axis).

(G) Table reports the p-value calculated from a Chi-squared test when the null hypothesis is to fit a combined data set between two pairs into a single population distribution, while the alternative hypothesis is to fit into a two-population distribution. Experimental conditions with significant difference ($p < 0.01$) are colored red.

A

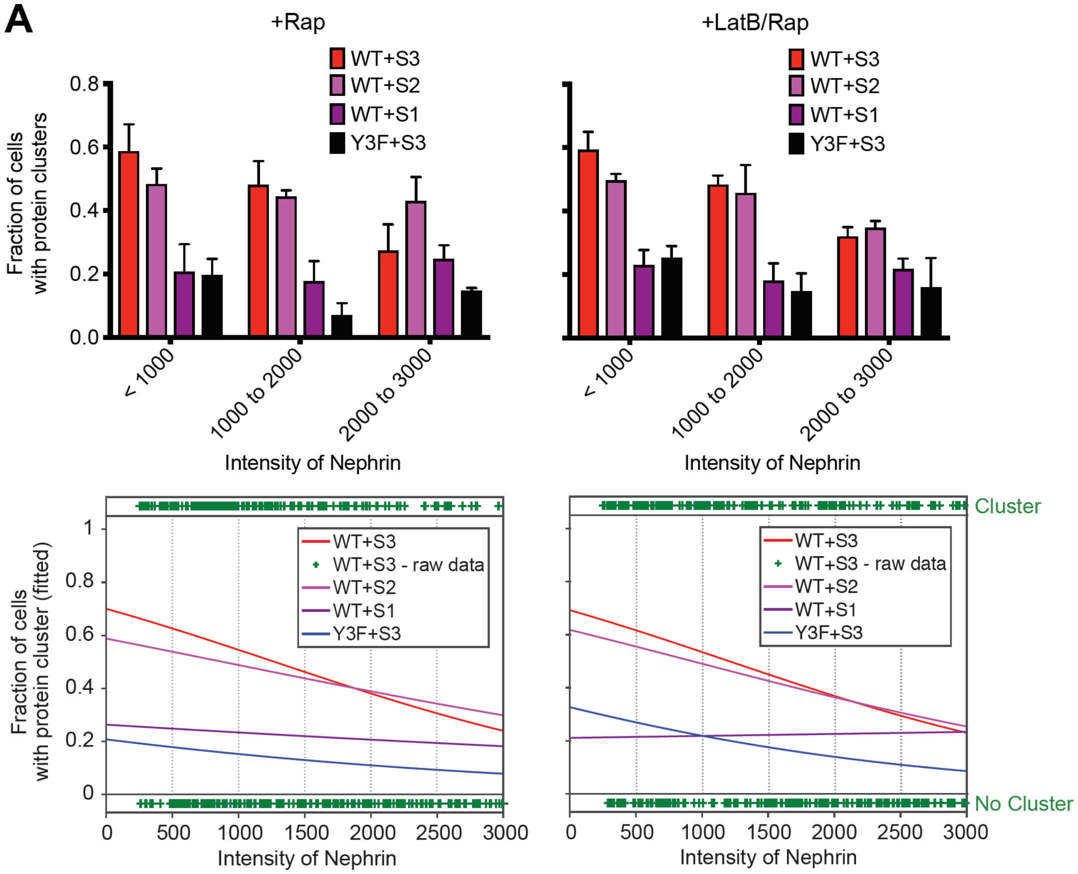

B

| | | Rap | | LatB/Rap | |
|---|---|---|---|---|---|
| | | Nephrin | Nck1 | Nephrin | Nck1 |
| WT + S3 | WT + S2 | 7e-1 | 8e-1 | 7e-1 | 1e-4 |
| WT + S3 | WT + S1 | <0.0001 | <0.0001 | <0.0001 | <0.0001 |
| WT + S3 | Y3F + S3 | <0.0001 | <0.0001 | <0.0001 | <0.0001 |
| WT + S2 | WT + S1 | <0.0001 | <0.0001 | <0.0001 | <0.0001 |
| WT + S2 | Y3F + S3 | <0.0001 | <0.0001 | <0.0001 | <0.0001 |
| WT + S1 | Y3F + S3 | <0.0001 | 7e-3 | 5e-4 | 4e-1 |

**Supplemental Figure S4. Nephrin/Nck1 clusters at the basal membrane are dynamic and their formation is dependent on valency.**

(A) Quantitative analyses of protein clustering with different valencies and cellular concentrations of Nck1. TIRF images of Nephrin and epi/TIRF images of Nck1 from fixed HeLa cells were used for the analysis. Cells expressed Src-FKBP, Nephrin(WT or Y3F as indicated)-FRB, and one of engineered Nck constructs, S3, S2 or S1. Graphs on the left were from cells treated with 10 min of MβCD followed by 30 min of rapamycin, while those on the right were from cells treated with 10 min of MβCD followed by 15 min of latrunculin B and then 15 min of rapamycin prior to fixation. (N = 3 independent experiments, each with > 125 cell images) Graphs on the top row show the fraction of cells with clusters for different membrane expression levels of Nephrin (determined from TIRF fluorescence intensity level of Nephrin). Error bars, s.e.m. Graphs on the bottom row show the probability of a cell forming clusters at a given Nephrin intensity level on the membrane. Cell data for each condition were fitted (lines) with a logistic regression model (left axis). Each plus sign indicates a status of a single cell (containing or lacking clusters) from 'WT + S3' sample (right axis).

(B) Table reports the p-value calculated from a Chi-squared test when the null hypothesis is to fit a combined data set between two pairs into a single population distribution, while the alternative hypothesis is to fit into a two-population distribution. Experimental conditions with significant difference ($p < 0.01$) are colored red.

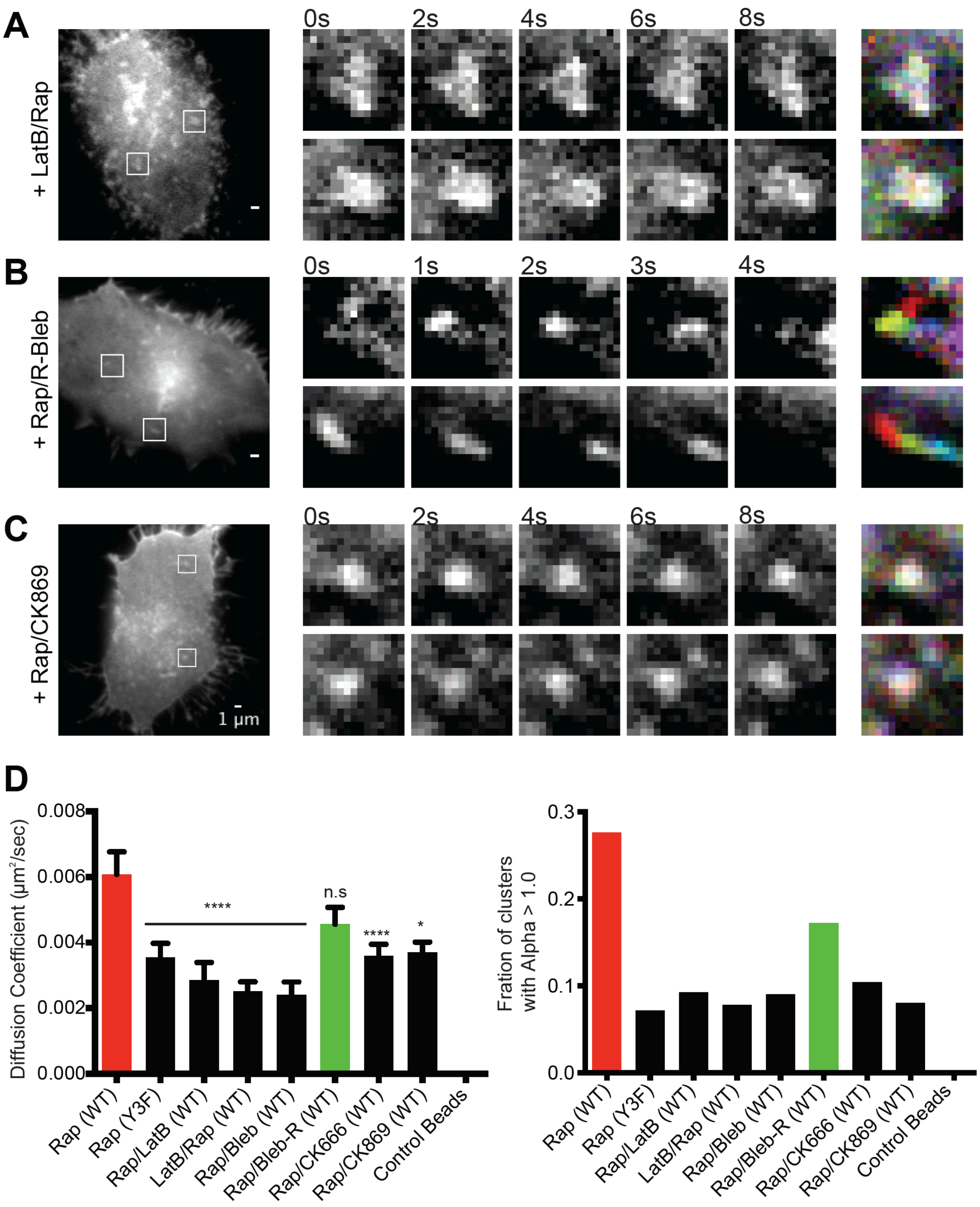

**Supplemental Figure S5. Movement of Nephrin clusters is dependent on the cortical actomyosin network.**

TIRF images for analysis were from live cells expressing Src-FKBP, Nephrin (WT or Y3F as indicated)-FRB (shown in images), and Nck1. Images were recorded every 250 ms for 30 s. Prior to imaging, cells were treated with 10 min of MβCD and 30 min of rapamycin (Rap) or 10 min of MβCD, 20 min of latrunculin B followed by 20 min of rapamycin (LatB/Rap). Alternatively, after 10 min of MβCD and 20 min of rapamycin, cells were treated with 20 min of latrunculin (Rap/LatB), active blebbistatin (Rap/Bleb), inactive blebbistatin (Rap/R-Bleb), CK666 (Rap/CK666), or CK869 (Rap/CK869).

(A to C) Timelapse images of cells treated with (A) LatB/Rap, (B) Rap/R-Bleb and (C) Rap/CK869. In the left images, two clusters are indicated with arrows in each cell. These clusters are shown in expansion in the time courses to the right. The right-most column shows overlays of the five timecourse images coded with different colors. In this representation, when movement is limited, clusters at different time points superimpose and appear white. Contrast and brightness settings for the different cells were individually optimized to aid visualization of clusters.

(D and E) Error bars, s.e.m. $N \geq 14$ cells counted for each condition. (*$p < 0.05$, ****$p < 0.0001$)

(D) Average diffusion coefficient of clusters in cells treated with actomyosin inhibitors.

(E) Fraction of cells with calculated alpha > 1.0 from time-lapse images of cells treated with actomyosin inhibitors.

**A**

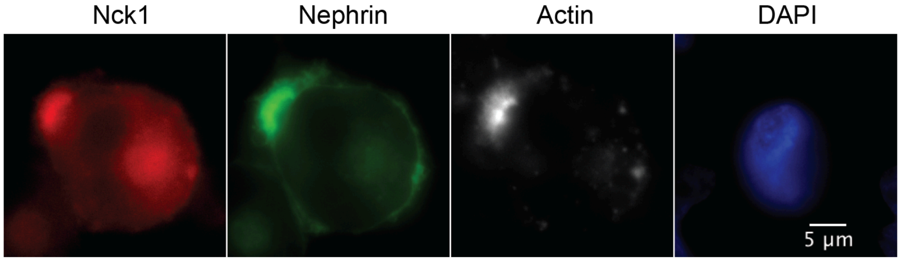

**B**

| | Nephrin | | | | |
|---|---|---|---|---|---|
| | DMSO | Rap (clustered) | Rap (unclustered) | Rap/LatB (clustered) | Rap/LatB (unclustered) |
| Preferred model | Two phase decay | Two phase decay | Two phase decay | Two phase decay | Two phase decay |
| Percent Recovery | 97 | 73 | 88 | 58 | 10 |
| PercentFast | 49 | 30 | 33 | 22 | 57 |
| Half Life (Slow) | 39 | 30 | 9 | 29 | 44 |
| Half Life (Fast) | 4 | 3 | 2 | 2 | 5 |

| | Nck1 | | | | |
|---|---|---|---|---|---|
| | | Rap (clustered) | Rap (unclustered) | Rap/LatB (clustered) | Rap/LatB (unclustered) |
| Preferred model | | Two phase decay | Two phase decay | Two phase decay | Two phase decay |
| Percent Recovery | | 79 | 104 | 76 | 102 |
| PercentFast | | 40 | 66 | 19 | 46 |
| Half Life (Slow) | | 22 | 24 | 14 | 6 |
| Half Life (Fast) | | 3 | 2 | 1 | 1 |

| | N-WASP | | | | |
|---|---|---|---|---|---|
| | | Rap (clustered) | Rap (unclustered) | Rap/LatB (clustered) | Rap/LatB (unclustered) |
| Preferred model | | Two phase decay | Two phase decay | Two phase decay | One phase decay |
| Percent Recovery | | 88 | 100 | 51 | 98 |
| PercentFast | | 39 | 60 | 40 | N/S (single decay) |
| Half Life (Slow) | | 28 | 26 | 31 | N/S (single decay) |
| Half Life (Fast) | | 4 | 2 | 7 | 9 |

**Supplemental Figure S6. Nephrin/Nck1 clusters at the cell periphery are resistant to latrunculin B treatment.**

(A) Epi fluorescence images of a fixed cell expressing Src-FKBP (not shown), Nephrin-FRB (green), Nck1 (red). Cells were stained for F-actin (magenta) with A647-phalloidin and for nucleus (blue) with DAPI. Cells were treated with 10 min of MβCD, 10 min of rapamycin followed by 10 min of latrunculin B.

(B) Tables show the parameters from fitting the FRAP data for Nephrin (top), Nck1 (middle), and N-WASP (bottom). F-tests were performed to determine whether the data were best fit to single or double exponential decay functions. $N > 25$ cells analyzed for Nephrin. $N \geq 14$ cells analyzed for Nck1 and N-WASP.
